# ACTIVITY IN HUMAN DORSAL RAPHE NUCLEUS SIGNALS CHANGES IN BEHAVIOURAL POLICY

**DOI:** 10.1101/2025.01.08.632066

**Authors:** Luke Priestley, Ali Mahmoodi, William Reith, Nima Khalighinejad, Matthew Rushworth

## Abstract

The dorsal raphe nucleus (DRN) is an important source of serotonin to the human forebrain, however there is little consensus about its behavioural function. We build on recent results from animal models to demonstrate that activity in human DRN represents changes between general behavioural policies. We use a novel behavioural task to show that human participants change their policy to pursue or reject reward opportunities as a function of the average value of opportunities in the environment. Activity in DRN – but no other neuromodulatory nucleus – signalled such policy changes. Patterns of multivariate activity in dorsal anterior cingulate cortex (dACC) and anterior insular cortex (AI), meanwhile, tracked the relative value of reward opportunities given the average value of the environment. We therefore suggest that DRN, dACC and AI form a circuit in which dACC/AI compute the relative value of reward opportunities given the current context, and DRN implements changes in behavioural policy based on context-specific values.

## Introduction

What makes an opportunity worth pursuing? For both humans and non-human animals, these judgments depend not just on the reward itself but on the context in which the judgements are made. A quintessential example is the prey selection dilemma set forth in behavioural ecology, in which a predator must decide whether to seize a foreground opportunity available in the present or search the environment for better alternatives (Charnov, 1976; Krebs et al., 1977). According to normative accounts of this scenario, an opportunity should only be pursued if its value meets or exceeds the opportunity cost of obtaining it. In practice, this means that opportunities with the same value will evoke different behavioural policies depending on the context they arise in: for example, it is rational to pursue even an objectively mediocre opportunity in a poor environment where everything on the horizon is similar or worse, and yet the same opportunity might be rejected in the context of a rich environment with plentiful high-value alternatives.

This model argues that organisms should compare any opportunity they encounter with the potential alternative opportunities they would forgo if they engage with it. While organisms are unlikely to have precise knowledge of all potential future opportunities, previous opportunities constitute a proxy for future opportunities. Therefore, the computational problem for the brain is to track recent opportunities (is the environment on average a rich one or poor one) and to adjust the policy for rejecting or accepting each type of opportunity that might be encountered. As a result, whether a middling opportunity should be accepted or rejected should depend on whether the average value of the environment has recently been poor or rich.

How does the brain adapt behavioural policies to the environment in this way? In this study, we argue for the importance of a cortico-subcortical circuit comprised by dorsal raphe nucleus (DRN) – a brainstem nucleus that is the primary source of serotonin to the forebrain in mammals – and anterior cingulate cortex (ACC) and anterior insular cortex (AI) (Charara & Parent, 1998; Hornung, 2003; Huang et al., 2019). Although serotonin has attracted broad interest – partly due to its link with affective disorders –, little is known about what DRN activity represents, and how DRN function differs from other neuromodulatory nuclei in the brainstem, midbrain, and basal forebrain. Our approach builds on previous studies in both human and non-human primates showing that activity in ACC and AI tracks reward-related statistics that are critical for adaptive decision-making, like the value, valence and background-rate of recent reward outcomes (Hayden et al., 2011; Khalighinejad et al., 2021; Kolling et al., 2012; Wittmann et al., 2020). Similarly, it builds upon a rich vein of experiments in animal models linking DRN activity to learning and decision-making. A consistent theme in these studies is that DRN controls the balance between opponent modes of reward-driven behaviour, like patience and impulsivity, exploration and exploitation, and even motivation and demotivation for reward-seeking (Khalighinejad et al., 2022; Lottem et al., 2018; K. W. Miyazaki et al., 2014; Priestley et al., 2024). Here, we argue that these seemingly disparate functions involve a common process of setting a behavioural policy. We employ a novel behavioural task that allows us to identify when and how behavioural policies are set, and demonstrate for the first time that activity in human DRN is associated with policy changes.

We gave participants a novel, foraging-inspired behavioural task in which they encountered a limited set of reward options in a series of rich and poor environments. As expected, participants systematically changed their behavioural policy depending on the richness of the environment, and were especially responsive to poor environments which caused them to become less selective in their choices. We recorded brain activity while participants performed the task using an ultra-high field functional magnetic resonance imaging (fMRI) protocol specifically designed to maximise signal from brainstem and midbrain areas. These recordings demonstrated that DRN – but no other major neuromodulatory nucleus – exhibited distinctive univariate patterns of activity that were specific to behavioural-policy changes. In contrast, multivariate patterns of activity evoked by reward-options in dACC and AI paralleled the relative-value of each option, as apparent in each participant’s behavioural policy, as the environmental context shifted._Taken together, these results suggest that DRN, dACC and AI form a cortico-subcortical circuit that reconciles the brain’s behavioural policy with the distribution of rewards in the environment.

## Results

### A behavioural task for manipulating the richness of the environment

Twenty-seven participants performed a simple decision-making task involving sequential encounters with reward opportunities (fig.1A). On each trial, participants were presented with a single reward opportunity drawn from a set of three possible options with known *a priori* values: a low-value option (5-points; bronze medallion), a middle-value option (10-points; silver medallion), and a high-value option (50 points; gold medallion). Upon each, encounter, participants could pursue opportunities by making a button press response or reject opportunities by withholding their response and waiting for the next trial to begin (see Methods).

**Figure 1.**
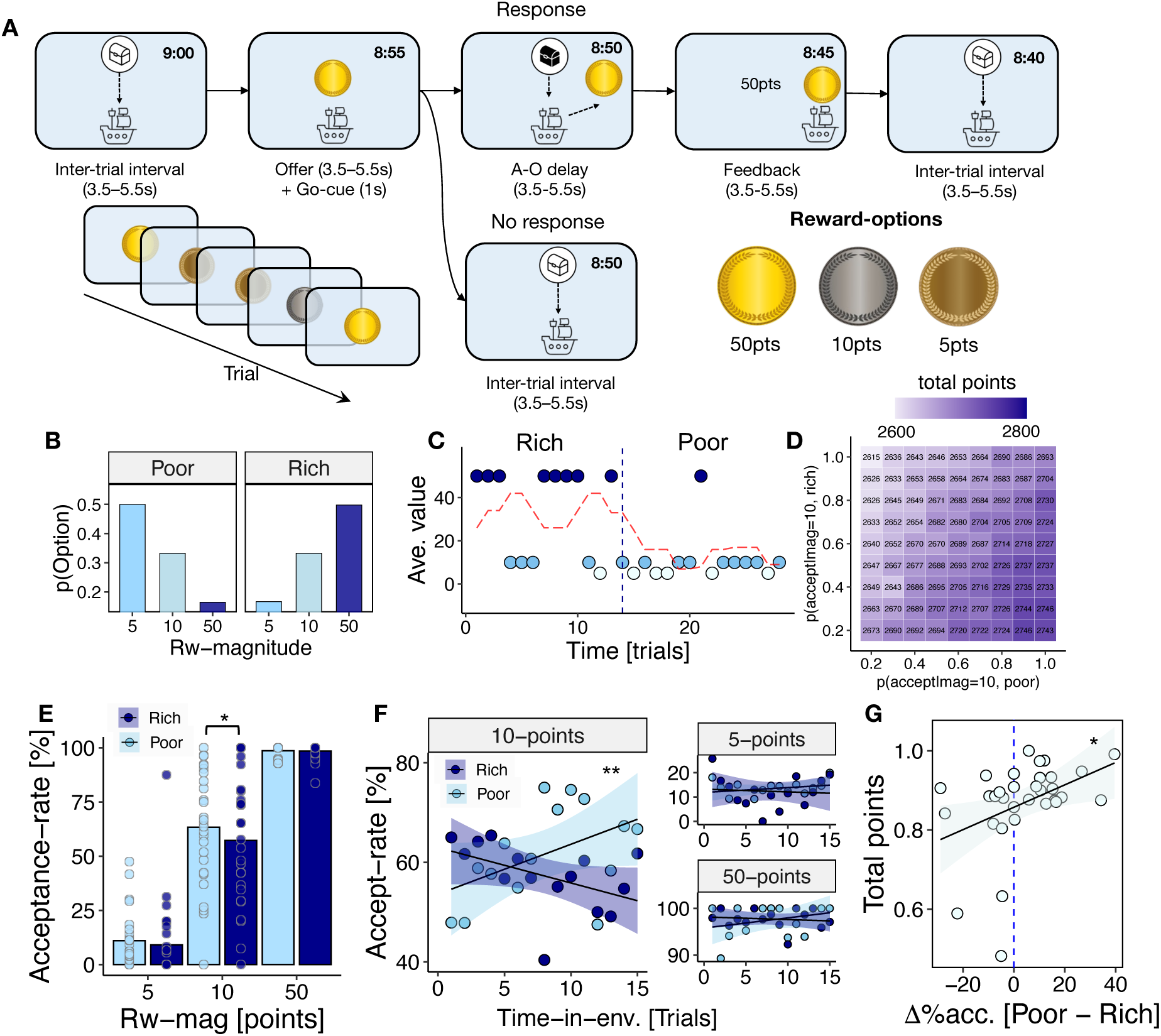
Reward-guided decisions are modulated by the richness of the environment. **(A)** Subjects played the role of a treasure seeking ship’s captain in a simple behavioural task involving sequential encounters with reward-options. A single reward-option was encountered on each trial, and subjects needed to decide whether to pursue (button-press response) or reject (no response) that option. Pursuing an option incurred a temporal opportunity cost that was equivalent to giving up a future encounter, which was symbolised by a passing black treasure-chest during the action-outcome delay period. To earn as many points as possible, subjects therefore needed to compare the reward-option available on each trial with the distribution of opportunities they expected in the future. Reward-options were symbolised by gold (50-points), silver (10-points) and bronze (5-points) medallions. The remaining time in the session was shown on the top left corner of the screen. **(B)** The frequency of 5- point, 10-point and 50-point reward-options was systematically manipulated to engender differences in the richness of the environment. In rich blocks, the 50-point option was more frequent than 10-point and 5-point options, whereas in poor blocks the 5-point option was more frequent than 10-point and 50-point options. Note that the frequency of the 10-point option remained constant, meaning that behaviour toward the 10-point option could be compared across blocks without bias. **(C)** An example sequence of rich and poor blocks illustrating key reward statistics and their evolution over time. Coloured dots indicate the specific reward-option available on each trial (dark-blue = 50-points, mid-blue = 10-points, light-blue = 5-points). Dashed red line indicates the mean value of rewards over the preceding five trials (i.e., *t*-1:*t*- 5). **(D)** The temporal and reward-value parameters of the task were carefully chosen to engender changes in behaviour toward the 10-point option. Tile-map and colour- scale shows total points earned by simulated agent performing the experiment as a function of pursue-rates for the 10-point option in rich-environments (y-axis) and poor environments (x-axis). Note that environments were not signalled to participants, and so we did not expect them to display this pattern of behaviour immediately upon entering a new environment. Instead, we expected that they would approach the pattern as they accumulated experience within an environment. **(E)** Subjects pursue the 10-point option more frequently in poor-environments relative to rich environments. In contrast, there are no behavioural differences for 5-point and 50- point environments as a function of environment-type. Bars and dots indicate sample-level and subject-level mean pursue-rates, respectively, in each condition. **(F)** Environments were not explicitly signalled to subjects and needed to be inferred from the relative frequency of reward-options over time. Accordingly, the difference in middle-option pursue-rates (y-axis) for poor-environments and rich-environments increased as a function of time spent in an environment (x-axis). There were no such changes for 5-point and 50-point options. Dots and whiskers indicate mean ± SEM of pursue-rate in each level of trial-within-environment (x-axis). **(G)** The degree to which subjects changed their behaviour between environments predicted their success on the task. Y-axis denotes total-points earned in the experiment and x-axis denotes mean difference in middle-option pursue-rates between poor-environments and rich-environments [poor – rich]. Dots represent data from individual subjects.

Pursue decisions incurred a temporal opportunity cost equivalent to foregoing a single reward opportunity in the future. It was therefore critical to consider the prospective frequency of each reward-option when deciding whether an opportunity was worth pursuing. The frequency of reward-options was systematically manipulated by dividing the task into 4.5-minute blocks of trials with different reward-option distributions (fig.1B). There were two types of block; (1) *rich* blocks, where the 50-point option was more frequent than the 10-point and 5-point options *P(5- point|Rich)* = 0.16, *P(10-point|Rich)* = 0.33, *P(50-point|Rich)* = 0.50), and; (2) *poor* blocks where the 5-point option was more frequent than the 10-point and 5-point options (*P(5-point|Poor)* = 0.50, *P(10-point|Poor)* = 0.33, *P(50-point|Poor)* = 0.16). These blocks allowed us to experimentally control the richness of the environment throughout the task (fig.1C). Importantly, the frequency of the 10-point option was constant across both rich and poor blocks, meaning that changes in a participant’s strategy in relation to the 10-point option could only arise from changes in the environmental context within which it occurred.

The parameters of the task were carefully chosen to ensure that the rational task-strategy was to accept the 10-point option in poor environments, and to reject it in rich environments (fig.1D). Environments were not signalled to participants, and we therefore did not expect participants to exhibit precisely this pattern. Nevertheless, the design ensured that there was a reward-maximising rationale for participants to change their behaviour toward the 10-point option. Accordingly, we focus predominantly on 10-point option trials in the following behavioural analyses, although the same conclusions obtain in omnibus analyses where all reward-options are considered together (GLM1.1a; see supplementary fig.S1).

#### Reward-guided decisions are modulated by the richness of the environment

To begin, we tested the hypothesis that participants were more likely to pursue the 10-point option in poor environments compared to rich environments. All analyses in this section were implemented using mixed-effect generalised linear models (GLM) which account for subject-specific variation in behaviour, and in which the dependent variable was the binary pursue-vs-reject decision on each trial.

We first tested the hypothesis by asking whether the experimentally manipulated environment-type (rich-vs-poor) influenced pursue-vs-reject decisions. This showed that participants were, indeed, more likely to pursue the 10-point option when it occurred in a poor environment compared to a rich environment (GLM1.1b; β_environment-type_ = -0.19, SE = 0.10, *p* = .045; fig.1E). There was no evidence for similar effects in 5-point (GLM1.1b; β_environment-type_ = -0.58, SE = 0.31, *p* = .062; fig.1E) or 50- point options (GLM1.1b; β_environment-type_ = 0.64, SE = 0.51, *p* = .206; fig.1E). Analysing all options together in a single GLM indicated a two-way interaction between option-value and environment-type that was consistent with the pattern evinced by option-specific analyses (GLM1.1a β_environment-type*reward-option_ = -3.09, SE = 0.19, *p* < .001). Importantly, the difference in 10-point option pursue-rates between poor and rich environments [Δ(pursue-rate) = pursue-rate_poor_ – pursue-rate_rich_] was positively correlated with total points earned in the experiment (*r_Δ(pursue-rate) vs total-points_* = 0.40, *p* = .040; fig.1G). The degree to which participants changed their behaviour between environments thus determined their performance on the task, as envisaged in the experimental design.

Participants were not informed that there were distinct rich and poor environments, and they received no cues to this effect during the task. Environment-driven changes in behaviour therefore involved tracking the frequency of reward- options over time. To investigate whether the environment effect emerged over time in this way, we tested the two-way interaction between environment-type and time elapsed within an environment, where time was quantified in trials. This revealed that environment-driven changes in behaviour increased as a function of time for the 10- point option (GLM1.2 β_environment-type*trial-in-environment_= -0.20, SE = 0.06, *p* = .001; fig.1F), but that there were no such changes for the 5-point (GLM1.3b; β_environment-type*trial-in-environment_=-0.04, SE = 0.12, *p* = .693; fig.1F) and 50-point options (GLM1.2; β_environment-type*trial-in- environment_= -0.43, SE = 0.25, *p* = .092; fig.1F). Simple main-effects analysis for the 10- point option indicated an asymmetric pattern of responsiveness to environments, whereby participants were more likely to pursue the 10-point option as time elapsed in poor environments (GLM1.2; β_trial-in-environment_= 0.27, SE = 0.08, *p* < .001), but were not more likely to reject the 10-point option over time in rich environments (GLM1.2; β_trial-in-environment_= -0.13, SE = 0.09, *p* = .165). This pattern of results was confirmed by a conceptually similar analysis in which the environment was operationalised as the average value of recently encountered reward-options, as distinct from the experimentally defined environments (GLM1.3; see supplementary fig.S1C-D).

#### Option-specific policies are reconciled with the richness of the environment

Having shown that pursue-vs-reject decisions were influenced by the richness of the environment, we performed a series of complementary analyses focussed on the temporal organisation of behaviour. Because the task featured only three reward- options, we reasoned that participants would form a behavioural policy for each option – that is, an option-specific pursue-vs-reject strategy that remained consistent over successive encounters with the option in question (fig.2A). We validated this hypothesis by testing whether pursue-vs-reject decisions were autocorrelated across encounters with the same reward-option. This confirmed that participants tended to repeat their decisions across successive encounters, consistent with a temporally persistent behavioural policy (GLM2.1; β_previous-policy_ = 1.53, SE = 0.28, *p* < .001; fig.2B). Importantly, when the pursue-vs-reject decision on the preceding trial concerned a different option, it did not influence the pursue-reject decision on the current trial, suggesting that policies were option-specific (GLM2.2; β_previous-action_= 0.05, SE = 0.17, *p* =.737; fig.2B; see also fig.2A for visual explanation of distinction between switching in relation to a previous-action and switching in relation to a previous-policy).

**Figure 2.**
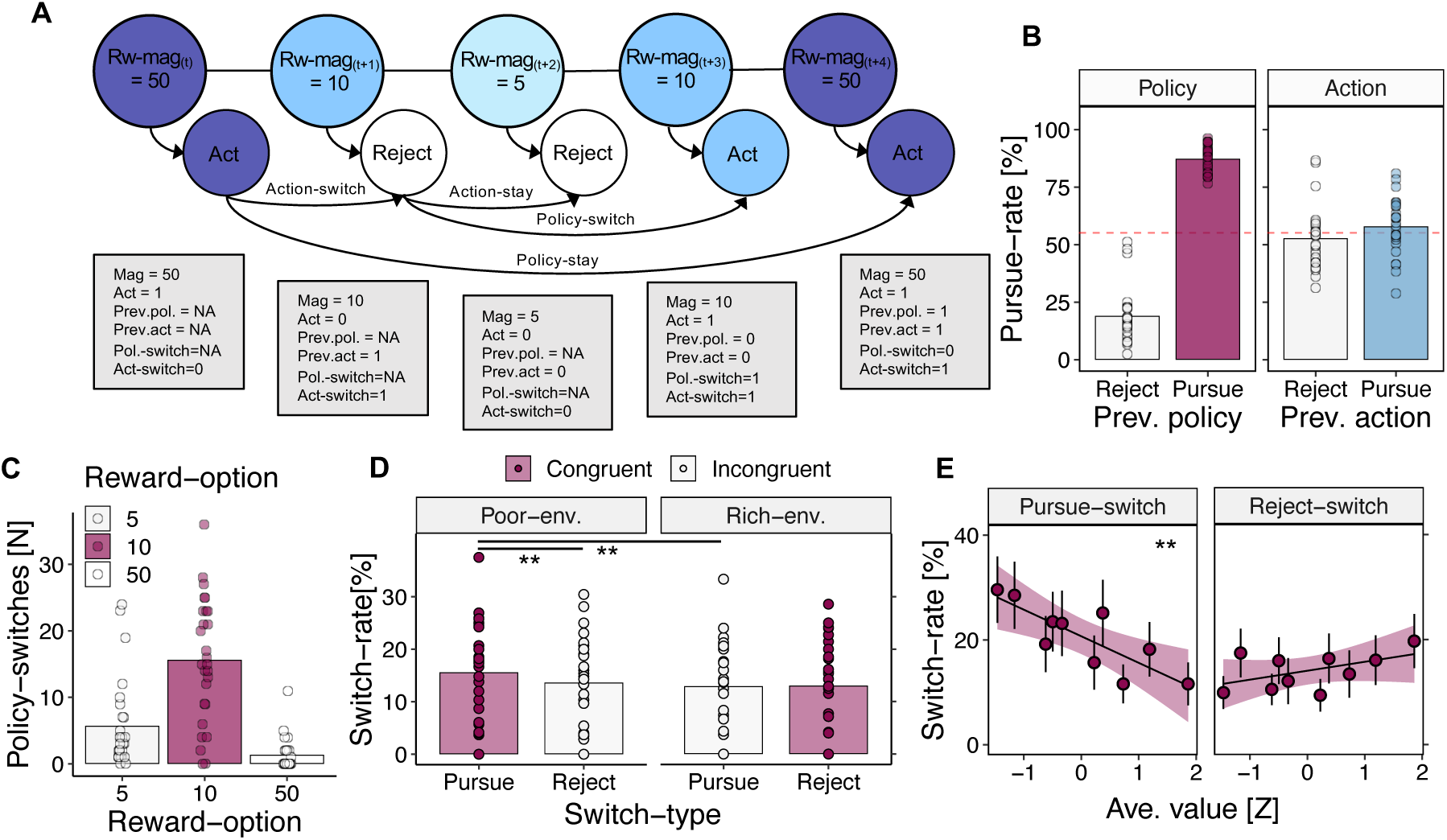
Subjects exhibit consistent behavioural policies that are shaped by the richness of the environment. **(A)** Employing a limited set of reward-options in the task allowed us to precisely track changes in behaviour. We categorised these changes along two dimensions: (1) a *policy* dimension reflecting the option-specific decision to pursue-vs-reject each of the three available reward-options, and (2) an *action* dimension reflecting the non-option-specific decision to pursue-vs-reject opportunities regardless of the reward-option available. Within this framework, we then categorised trials as (i) *policy-stay* events if the pursue-vs-reject decision was consistent with the previous encounter, or (ii) *policy-switch* events if the pursue-vs- reject decision was different to the previous encounter. Correspondingly, we defined trials as (i) *action-stay* if the pursue-vs-reject decision was the same as the immediately preceding trial (regardless of reward-option), and (ii) *action-switch* if the pursue-vs-reject decision changed. **(B)** Participants were more likely to pursue a reward-option if they pursued it on the previous encounter and vice versa (previous- policy effect; left panel). Conversely, pursue-vs-reject decisions were not influenced by the action made on the previous trial (previous-action effect; right panel). Bars show sample-mean pursue-rate in each condition. Dots indicate subject-specific pursue-rates. Dashed red line indicates overall mean pursue-rate, regardless of previous-policy and/or previous-action condition. **(C)** Policy-switches were more frequent for the 10-point option than for 5-point and 50-point options. Bars show overall mean count of policy-switches per experimental session. Dots indicate subject-specific counts. **(D)** Pursue-switches (i.e. switching from reject-to-pursue) are more likely to occur than reject-switches in poor environments, consistent with the rational strategy for performing the task. There was no corresponding difference between pursue-switches and reject-switches in the rich-environments. Colour-scale denotes congruency of switch-direction relative to environment-type – i.e., the switch-direction that subjects should display if following the rational strategy. Data shown for 10-point option only. Bars indicate overall mean rate of switch-events and dots indicate subject-specific rates of switch events. **(E)** Probability of pursue-switch (left) and reject-switch (right) events when the richness of the environment is operationalised as the average value of recent reward-options. Similar to (D), pursue- switches are more likely to occur as the richness of the environment decreases (i.e. in poor environments), but reject-switches follow no systematic pattern. Data shown for 10-point option only. Points and error bars indicate mean ± SEM switch-rates in decile bins of average-value.

Adopting the behavioural-policy perspective enabled us to precisely identify changes in behavioural policy (policy-switch, hence). We operationalised policy- switches as trials on which the pursue-vs-reject decision diverged from the policy exhibited the previous time a reward-option was encountered (fig.2A). Consistent with earlier analyses showing behavioural changes for the 10-point option, policy- switches were more frequent for the 10-point option than both its 5-point (GLM2.3; β_middle-vs-low_ = 1.52, SE = 0.39, *p* < .001; fig.2C) and 50-point counterparts (GLM2.3; β_middle-vs-high_ = 3.63, SE = 0.67, *p* < .001; fig.2C). Subsequent analysis of policy-switches therefore focused on the 10-point option trials (see supplementary fig.S2 for analysis of 5-point and 50-point options).

Given our analysis of pursue-vs-reject decisions, we predicted that policy- switches would be influenced by the richness of the environment. To test this, we categorised policy-switches into two-types depending on the direction of change: (1) *pursue-switches*, when the reward-option was pursued on the current trial after being rejected on the previous encounter, and (2) *reject-switches*, when the reward-option was rejected on the current trial after being pursued on the previous encounter. Insofar as participants approximated the optimal task strategy (fig1.D), we expected that pursue-switches would be more likely in poor environments and reject changes would be more likely in rich environments.

When pursue-switches were analysed in relation to the experimentally defined environments, they followed the expected pattern: in poor environments, pursue- switches were more likely to occur than reject-switches (GLM2.4a; β_pursue-switch vs reject- switch_= 0.89, SE = 0.19, *p* <.001; fig.2D), and pursue-switches in poor environments were also more likely to occur than pursue-switches in rich environments (GLM2.5a; β_rich-vs-poor_= -0.36, SE = 0.12, *p* = .002). The likelihood of reject-switches, however, did not differ across environment types (GLM2.4b, β_pursue-switch vs reject-switch_= -0.22, SE = 0.20, *p* =.274, fig.2D; GLM2.5b, β_rich-vs-poor_= 0.07, SE = 0.12, *p* = .550). A similar pattern obtained when policy-switches were compared against the average-value of recent reward-options: pursue-switches were more likely as average-value decreased (GLM2.6a; β_ave.-value_= -0.76, SE = 0.14, *p* <.001; fig.2E), but there was a null effect for reject-switches which occurred at an approximately constant rate (GLM2.6b; β_ave.- value_= 0.13, SE = 0.15, *p* = .386; fig.2E; see supplementary fig.S2 for equivalent analysis of 5-point and 50-point options). In tandem with previous analyses, these results suggest that changes in behavioural policy approximated the rational task strategy and that this pattern was more pronounced in poor environments.

#### Dorsal raphe nucleus represents environment-driven policy switches

Participants performed the behavioural task while undergoing ultra-high field fMRI recordings of blood oxygen level dependent (BOLD) signal. We used these recordings to investigate brain activity that represented the task environment, and environment-driven changes in behaviour. We focused on *a priori* regions of interest (ROIs) comprising the ascending neuromodulatory systems (ANS): an assemblage of phylogenetically ancient nuclei in the basal forebrain, midbrain, and brainstem that release fundamental neuromodulatory chemicals via diffuse, whole-brain connections (Dayan, 2012; Stephenson-Jones et al., 2013, 2016). According to a prominent theory, these nuclei broadcast salient low-dimensional signals that orchestrate activity across multiple brain regions (Dayan, 2012). These properties make the ANS ideal candidates for implementing changes in behavioural policy.

We extracted BOLD signal from an anatomical ROI for the dorsal raphe nucleus (DRN), in addition to a combined ROI covering the substantia nigra and ventral tegmental area (midbrain dopaminergic nuclei; MBD), the locus coeruleus (LC), the medial septal nucleus (MS), and the habenula – an epithalamic nucleus characterised by reciprocal interactions with the nuclei comprising the ANS ((Hikosaka, 2010; Matsumoto & Hikosaka, 2007); see fig.3A for anatomical illustration of all ROIs and Methods for details of their construction). Although our key hypotheses concerned DRN, expanding our purview allowed us to compare DRN activity with patterns in other ANS nuclei, which is critical for identifying the specialised function of different brain regions (Marshall et al., 2016).

**Figure 3.**
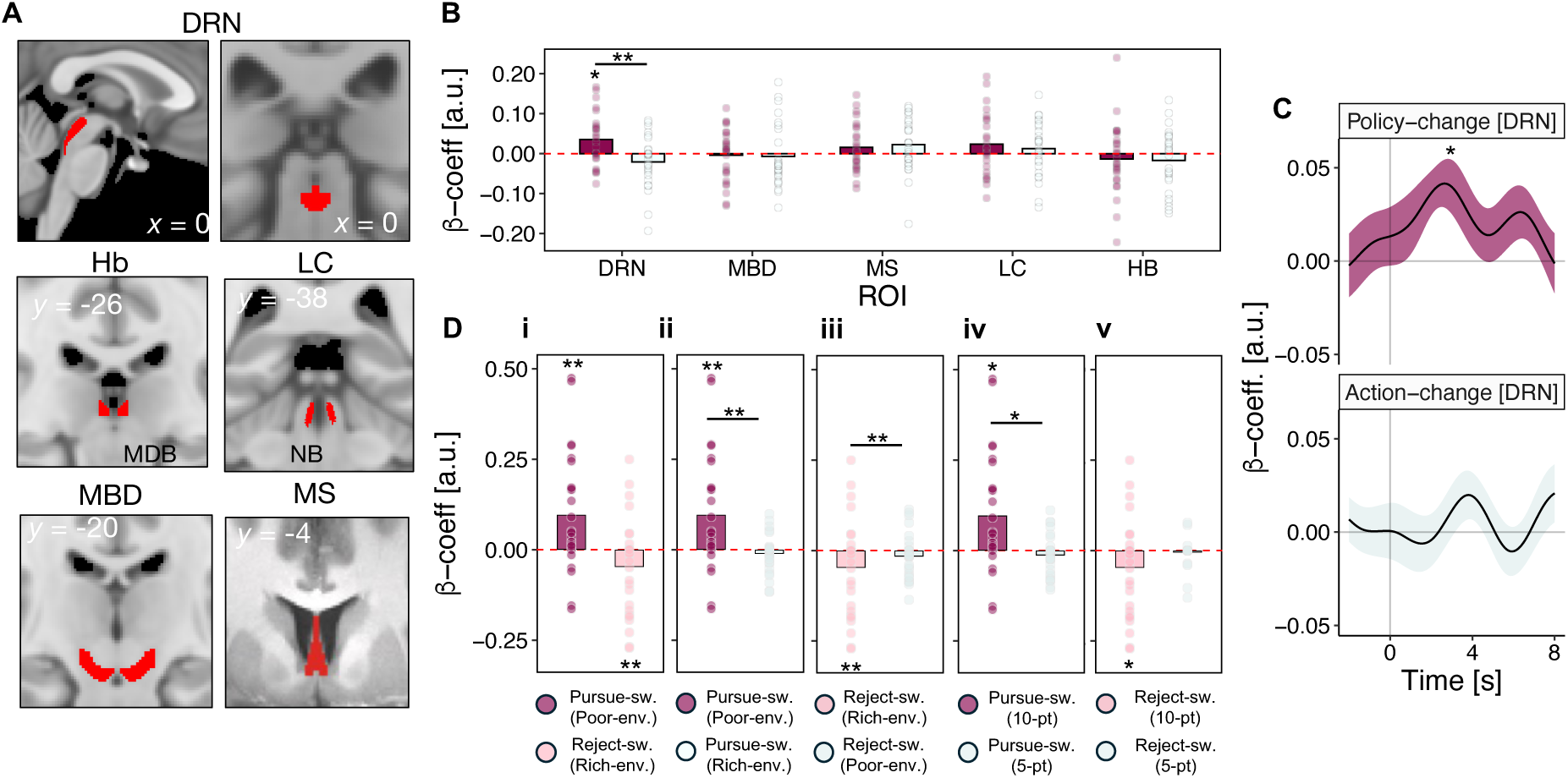
Brain activity in dorsal raphe nucleus represents the richness of the environment and environment-driven changes in behavioural policy. (A) The. fMRI analysis focussed on anatomical regions of interest (ROIs) in the ascending neuromodulatory systems (ANS). ROIs comprised the dorsal raphe nucleus (DRN), habenula (Hb), locus coeruleus (LC), substantia nigra and ventral tegmental area (MDB), and nucleus basalis (NB). See Methods for details of mask construction. **(B)** Distribution of peak regression-weight for policy-change (purple) and action-change (grey) events in subcortical ROIs. Policy-changes – but not action-changes – modulate DRN activity, suggesting that DRN is involved in option-specific forms of behavioural change. No other ROIs showed representations of behavioural change. **(C)** Timecourse of the relationship between BOLD signal and: (i) policy-change events (top; purple), and; (ii) action-change events (bottom; grey) in DRN. Time=0 (x-axis) corresponds to onset of reward-opportunity stimulus. Given haemodynamic delay, it is likely that policy-change related neural activity in DRN occurs in ITI periods, when subjects are integrating reward feedback from the previous trial and planning their future behaviour. **(D)** Peak regression-weights for control analyses linking DRN to environment-driven policy switches: **(i)** shows DRN activity during policy changes that are congruent to the environment – i.e., pursue-change events in poor environments (dark-pink) and reject-change events in rich environments (light-pink); **(ii)** shows DRN activity during pursue-change events in poor environments (dark-pink) and rich- environments (grey) – i.e., events matched on motor output, but that differ in environment-congruence; **(iii)** shows DRN activity during reject-change events in rich environments (light-pink) and poor-environments (grey) – i.e., events matched on motor output, but that differ in environment-congruence; **(iv)** shows DRN activity during pursue-change events for the 10-point option (dark-pink) and 5-point option (grey), where only policy-switches for the 10-point were modulated by the environment in behaviour (fig.2C–D; supplementary fig.S2); **(v)** shows DRN activity during reject-change events for the 10-point option (light-pink) and 5-point option (grey). In timecourse graphs (C), lines and shadings show the mean and standard error (SE) of the β weights across participants. In effect-size graphs (B, D), bars show sample-mean magnitude of the peak regression weight in the timecourse according to an unbiased leave-one-out procedure (see methods). Dots indicate peak regression weights for individual participants.

We first examined brain activity related to policy-switches. We did this by contrasting brain activity on policy-switch trials with brain activity on policy-stay trials – i.e., we asked whether specific patterns of brain activity occurred when a policy- switch was exhibited in behaviour. This indicated that activity in DRN – but no other subcortical ROI – represented policy-switch trials (GLM4.2; *t_DRN; policy-change_*(26)*=*2.84, *p*=.043; fig.3D-E). We confirmed that this effect was specific to behavioural-policy changes by testing DRN’s relationship with other forms of behavioural change, like action-switches where both the pursue-vs-reject decision and option-identity changed across consecutive trials such that the change in behaviour did not constitute change in option-specific policy (fig.2A). Action-switches had a null effect on DRN activity (GLM4.2; *t_DRN; action-change_*(26)*=*-1.69, *p*=.110; fig.3D–E), and the policy- switch representation reported earlier was, moreover, significantly different to the null action-switch signal (*t_DRN; action-switch vs policy-change_*(26)*=*3.18, *p*=.002; fig.3D–E).

Our analysis of behaviour showed that policy-switches were driven by the richness of the environment (fig.2D-E). Because brain activity in DRN represented policy-switches in general, we therefore next tested whether DRN represented environment-driven policy-switches specifically. We examined DRN activity during policy-switches for the 10-point option that were congruent with the environment – i.e., pursue-switches in poor environments, and reject-switches in rich environments. DRN activity correlated with both congruent pursue-switches and congruent reject- switches, albeit with oppositely signed patterns of activity (GLM4.3a, *t_DRN; pursue- switch_*(26)*=*2.79, *p*=.009; GLM4.3c *t_DRN; reject-switch_*(26)*=-*2.81, *p*=.009; fig.3F-i). Because pursue-switches were correlated with action and reject-switches were correlated with inaction, we ensured that these effects were not artefacts of motor preparation by comparing DRN activity for congruent-vs-incongruent pursue-changes, and congruent-vs-incongruent reject-switches: in other words, we compared events with the same motor-output that occurred in different environmental contexts. Unlike congruent-pursue changes (GLM4.3a, *t_DRN; congruent-pursue_*(26)*=* 2.79, *p*=.009; fig.3F-ii) incongruent-pursue changes (GLM4.3b, *t_DRN; incongruent-pursue_*(26)*=*-1.10, *p*=.280; fig.3F-ii) had a null effect on DRN activity, and the congruent pursue-switch signal was stronger than its incongruent-pursue counterpart (*t_DRN; congruent-vs-incongruent pursue_*(26)*=*2.60, *p*=.014; fig.3F-ii). Similarly, incongruent-reject changes had a null effect on DRN activity (GLM4.3d, *t_DRN; incongruent reject_*(26)*=-*1.35, *p*=.189; fig.3F-iii) in contrast to the significant effect of congruent-reject changes (GLM4.3c, *t_DRN; congruent reject_*(26)*=-*2.81, *p*=.009; fig.3F-iii), although the difference between these patterns was not significant (*t_DRN; congruent-vs-incongruent reject_*(26)*=-*1.30, *p*=.204; fig.3F-iii). In this way, DRN policy-switch representations paralleled behavioural adjustment to the environment (fig.1E-F): they were only significant during switches to pursuit and switches to rejection that were appropriate for the richness of the environment, and they were especially prominent for pursue-switches engendered by poor environments (fig.2D).

As a further test of DRN policy-switch signals, we compared congruent policy- switches for 10-point and 5-point options. Behavioural analysis (fig.2D–E and supplementary fig.S2) demonstrated that policy-switches for the 10-point option were modulated by the richness of the environment, whereas policy-switches for the 5-point option were not and instead appeared to be random or exploratory in nature. Comparing the two events therefore allowed us to isolate policy-switches driven by the richness of the environment from behavioural switches with other origins. Unlike the significant effect of congruent 10-point option pursue-switches (GLM4.3a, *t_DRN; 10- pt pursue-switch_*(26)*=*2.38, *p*=.024; fig.3F-iv), 5-point option pursue-switches in poor environments had a null effect on DRN activity (GLM4.3a, *t_DRN; 5-pt pursue-switch_*(26)*=*-1.23, *p*=.229; fig.3F-iv). These signals were significantly different from one another (*t_DRN; low- vs-middle pursue_* (26)*=*2.52, *p*=.020; fig. 3F-iv). Similarly, reject-switches for the 5-point option in rich environments had a null effect on DRN activity (GLM4.3c, *t_DRN; low-option reject_*(26)*=-*0.73, *p*=.471; fig.3F-v) in contrast to the significant effect of middle-option reject-switches (GLM4.3c, *t_DRN; middle-option reject_*(26)*=-*2.08, *p*=.046; fig.3F-v) although the difference between signals in this case was not significant (*t_DRN; low-vs-middle reject_*(26)*=* 1.77, *p*=.090; fig.3F-v). This provides further evidence DRN activity was linked to specifically environment-driven policy-switches.

It is important to note that the absence of policy-switch related activity in other ANS nuclei was not due to insensitivity in the fMRI signal recorded from these regions. For example, we observed patterns of MDB activity that are strikingly consistent with its role in reward-guided behaviour in previous studies: (i) a pattern related to the pursue-vs-reject decision on each trial, reminiscent of a reward-guided action initiation signal (Guitart-Masip et al., 2014; Khalighinejad et al., 2020), and; (ii) a pattern conveying the value-difference between the current reward opportunity and the richness of the environment, akin to a reward prediction-error (Schultz, 2006) (see supplementary fig.S3). Similarly, we found that in contrast to the representation of environment-driven policy-changes in DRN activity, policy-changes that were not appropriate given the richness of the environment were represented in MS activity (see supplementary fig.S6). This is broadly compatible with the link between acetylcholine and forms of exploratory behaviour, in which an organism deviates from an established reward-maximising policy (Marshall et al., 2016; Yu & Dayan, 2005). Taken together, these analyses suggest that activity in DRN – but no other ANS nucleus – represented changes in behavioural policy that aligned behaviour with the environment. The policy-switch effect was specifically linked to environment-driven changes in behaviour and was consistently more prominent for policy- switches engendered by poor environments. In this way, DRN policy-switch representations parallelled behavioural sensitivity to poor environments (fig.3Fi-v & fig.2D–E).

### Cortical multivariate value representations are modulated by the richness of the environment

Having shown that DRN activity represented environment-driven policy- switches, we expanded our fMRI analysis to cortical areas that might represent the behavioural policies themselves within the task context. It has been shown that the brain constructs abstract representational spaces in which choices can be compared along multiple dimensions depending on current behavioural relevance (Knudsen and Wallis, Cell, 2021; Park et al., Nature Neuroscience, 2021; Nitsch et al., Nat.Comm 2024, Mahmoodi et al, Neuron 2023, 2024). Because our behavioural data indicated that participants adaptively changed their behavioural policy regarding the middle option depending on context, we asked whether the relative positions of the middle options, in the abstract representational space used by the brain, also changed with context. We hypothesised that in regions encoding the task space, the distance between neural representations of each specific reward-option would change in tandem with the decision policy, given each context (fig.4A). We tested this using representational similarity analysis (RSA) on a series of ROIs deriving from functional fMRI analyses (Kriegeskorte et al., 2008). First, we searched for relevant ROIs in our own data using whole-brain fMRI analysis testing for regions that coded: (i) the pursue-vs-reject decision made on each trial, and; (ii) the reward-magnitude of the opportunity on each trial (GLM3.1 & GLM3.2; see supplementary fig.S4). This identified clusters of activity related to the pursue-vs-reject contrast in anterior insular cortex (AI), dorsal anterior cingulate cortex (dACC), and lateral frontopolar cortex (lFPC), and a cluster in subgenual anterior cingulate cortex (sgACC) related to the reward-magnitude contrast (fig.4B; see supplementary fig.S4 and tables 1 and 2 for full results of whole-brain analyses; see Methods for details of ROI construction). In addition, we investigated ROIs in dorsomedial prefrontal cortex (dmPFC) and pregenual anterior cingulate cortex (pgACC) based on activations reported in previous studies ((Kable & Glimcher, 2007; Klein-Flügge et al., 2022); fig.4B).

**Figure 4.**
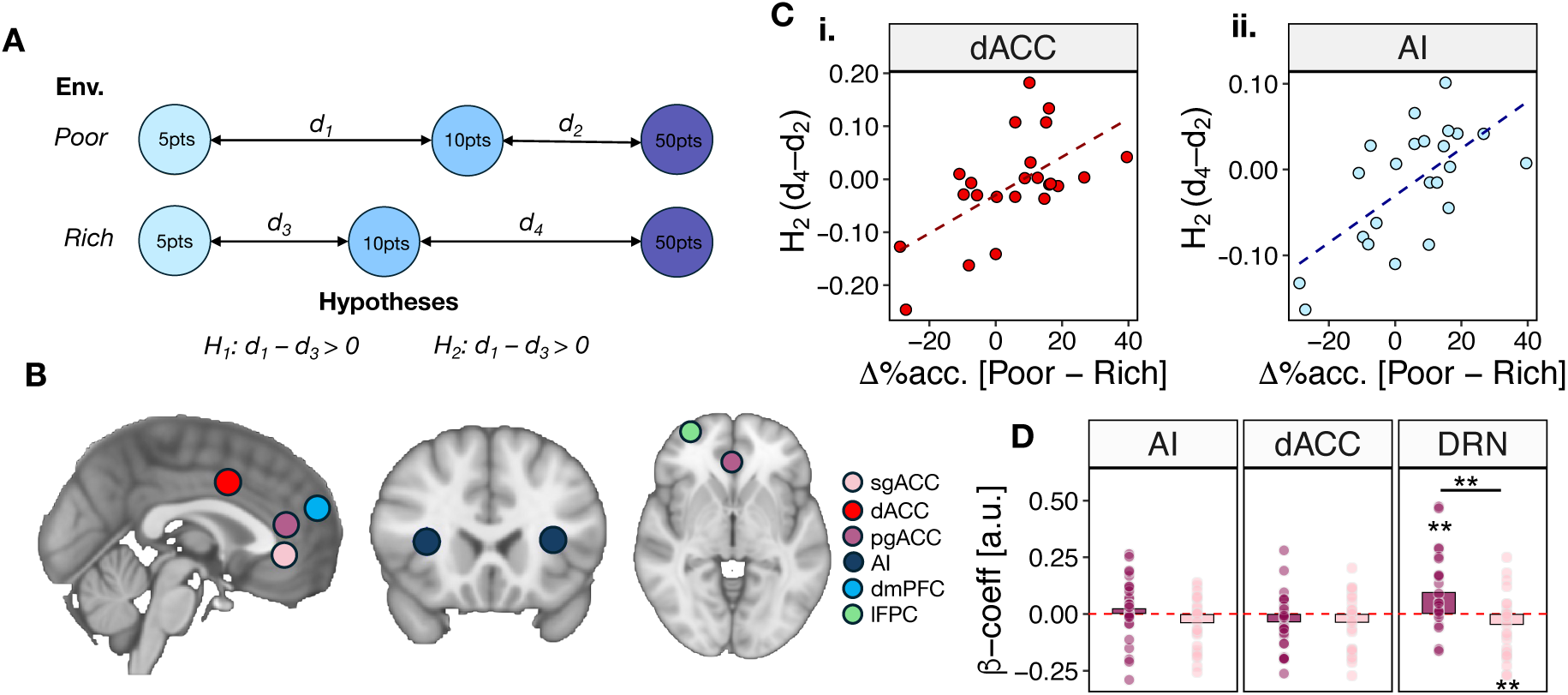
Multivariate representations of value are modulated by the environment in dorsal anterior cingulate cortex and anterior insular cortex. **(A)** We used representational similarity analysis (RSA) to examine how the distance between neural representations of reward options changed between rich and poor environments. Our hypothesis was that in regions encoding the task’s structure, the distance between representations would mirror changes in the relative value of the middle option. We therefore formulated two key hypotheses: (H_1_) that the 10-point option would be closer to the 5-point option in rich environments (d_3_) relative to poor environments (d_1_), i.e., d_3_ – d_1_ > 0, and (H_2_) that the 10-point option would be closer to the 50-point option in poor environments (d_4_) relative to rich environments (d_2_), i.e., d_4_ – d_2_ > 0. **(B)** We tested H_1_ and H_2_ in six cortical ROIs: subgenual anterior cingulate cortex (sgACC), dorsal anterior cingulate cortex (dACC), pregenual anterior cingulate cortex (pgACC), anterior insular cortex (AI), dorsomedial prefrontal cortex (dmPFC) and lateral frontopolar cortex (lFPC). Colour scale corresponds to ROI identity. **(C)** In dACC **(i)** and AI **(ii)**, d_4_ – d_2_ was positively correlated with the degree of behavioural change between environments – i.e., the more neurally similar the 10-point option was to the 50-point in poor environments, the more frequently subjects pursued the 10-point option in poor environments (and vice versa in rich environments). Points indicate individual participants. Dashed lines indicate line-of-best-fit. **(D)** In contrast to DRN (right-panel), univariate activity in dACC and AI did not covary with environment-driven changes in behavioural policy (see also fig.3). Bars show sample- mean magnitude of the peak regression weight according to an unbiased leave-one- out procedure (see methods). Dots indicate peak regression weights for individual participants. This suggests that although dACC and AI represent changes in the relative value of reward-options given the environment, they do not implement changes in behaviour that arise from those values (see also fig.5).

**Table 1:** fMRI cluster locations for GLM3.1

for $\beta_3$ decision(opportunity-onset) contrast in GLM 5.9
| Anatomical location | x | y | z | z-max | p | Size (# voxels) |
| --- | --- | --- | --- | --- | --- | --- |
| Left Thalamus | -15 | -21 | 7 | 5.28 | 9.61e-32 | 7132 |
| Postcentral Gyrus | -56 | -18 | 44 | 6.18 | 9.54e-31 | 6811 |
| Anterior Cingulate Gyrus | -1 | 5 | 38 | 4.95 | 1.6e-30 | 6739 |
| Right Cerebellum | 18 | -50 | -20 | 6.04 | 2.03e-29 | 6391 |
| Occipital Pole | 16 | -95 | 4 | 5.15 | 1.46e-19 | 3570 |
| Insular/Central Opperculum | -39 | -4 | 16 | 5.65 | 6.35e-16 | 2675 |
| Right Cerebellum | 17 | -58 | -46 | 4.72 | 1.58e-14 | 2355 |
| Left Cerebellum | -23 | -59 | -19 | 4.63 | 3.42e-11 | 1648 |
| Precentral Gyrus | -55 | 6 | 36 | 5.12 | 5.71e-11 | 1604 |
| Lingual Gyrus | -3 | -71 | 4 | 4.79 | 5.96e-08 | 1068 |
| Posterior Cingulate Gyrus/Precentral Gyrus | -8 | -24 | 44 | 5.16 | 8.34e-07 | 855 |
| Left Putamen | -32 | -8 | -2 | 4.44 | 7.45e-06 | 709 |
| Occipital Pole/Superior Lateral Occipital Cortex | -22 | -93 | 9 | 4.65 | 1.08e-05 | 685 |
| Inferior Lateral Occipital Cortex | -41 | -85 | 9 | 4.54 | 0.000365 | 471 |
| Lateral Frontal Pole (Right) | 30 | 59 | 0 | 4.14 | 0.000528 | 450 |
| Postcentral Gyrus/Anterior Supramarginal Gyrus | 40 | -31 | 41 | 4.31 | 0.00144 | 395 |
| Lateral Frontal Pole (Left) | -24 | 63 | 2 | 3.89 | 0.00155 | 391 |
| Insular/Central Opperculum (Right) | 44 | 2 | 11 | 3.91 | 0.00289 | 358 |
| Anterior Insular/Frontal Opperculum (Left) | -32 | 19 | 10 | 4.29 | 0.00461 | 334 |
| Cerebral White Matter (Right) | 24 | -23 | -4 | 4.56 | 0.00789 | 307 |
| Posterior Cingulate Gyrus/Precentral Gyrus (Right) | 16 | -31 | 39 | 4.01 | 0.0269 | 248 |
| Central Opperculum/Insular | 43 | 9 | 3 | 4.36 | 0.0334 | 238 |
| Anterior Insular/ Frontal Opperculum/ | 33 | 21 | 9 | 4.69 | 0.0484 | 221 |

**Table 2:** fMRI cluster locations for GLM3.2

for $\beta_3$ reward-magnitude contrast in GLM3.2
| Anatomical location | x | y | z | z-max | $p$ | Size (# voxels) |
| --- | --- | --- | --- | --- | --- | --- |
| Right Cerebellum | 20 | -50 | -20 | 4.35 | 9.48e-20 | 197 |
| Occipital Pole | 15 | -92 | 3 | 4.6 | 4.9e-15 | 135 |
| Postcentral Gyrus | -56 | -19 | 45 | 4.28 | 6.16e-13 | 110 |
| Insular/Central Opercular Cortex | -40 | -3 | 15 | 4.41 | 1.42e-10 | 84 |
| Postcentral/Supramarginal Gyri | -58 | -23 | 23 | 3.98 | 1.19e-07 | 55 |
| Lingual Gyrus | -4 | -74 | -2 | 4.15 | 1.31e-06 | 46 |
| Anterior Cingulate Cortex/Cingulate Gyrus | -6 | 6 | 43 | 4.05 | 5.13e-06 | 41 |
| Right Nucleus Accumbens | 6 | 11 | -5 | 4.71 | 2.12e-05 | 36 |
| Supramarginal Gyrus | -59 | -34 | 31 | 4.01 | 2.83e-05 | 35 |

Our analysis focussed on four representational distances; (i) *d_1_*, the distance between 5-point and 10-point options in poor environments; (ii) *d_2_*, the distance between 10 and 50-point options in poor environments; (iii) *d_3_*, the distance between 5-point and 10-point options in rich environments, and; (iv) *d_4_*, the distance between 10-point and 50-point options in rich environments. In each case, the distance was computed as the cosine dissimilarity between the pairs (1-cosine angle between the two offer values). Because this index is unaffected by changes in mean, univariate BOLD activity it should be unconfounded with differences in average brain activity associated with the contrasts that were first used to identify the ROIs. Moreover, note that the representational similarity analyses were time-locked to option presentation rather than to the later time of participants’ responses – the time point at which the ROIs were first identified. Insofar as the neural representation of reward- options tracked their relative positions in the task space given the environment, we predicted that distances would follow patterns where either: (1) H1: *d_1_* - *d_3_* > 0 – i.e., that the 10-point option’s representation was closer to the 5-point in rich environments compared to poor environments, or; (2) H2: *d_4_* - *d_2_* > 0 – i.e., that the 10-point option’s representation was closer to the 50-point option in poor environments compared to rich environments (fig.4A). Importantly, not all participants followed the rational strategy of pursuing the 10-point option more frequently in poor environments (fig.1E–F). We therefore anticipated that the magnitude of representational distance changes would be correlated with participant-specific levels of behavioural change between environments.

Comparing neural representations with behaviour demonstrated a striking effect in dACC and AI whereby the magnitude of H2 (*d_4_* - *d_2_*) was positively correlated with behavioural change between environments (*r_ACC_*=0.60, *t*(21)=3.46, *p*=.002; *r_AI_*=0.65, *t*(21)=3.90, *p*<.001; fig4C; correlation statistics reported after familywise error control using Bonferroni-Holm method) – in other words, in participants who adopted the rational strategy, neural representations of the 10-point option were closer to the 50-point option in poor environments than they were in rich environments. We confirmed this interpretation by contrasting *d_4_* - *d_2_* in participants who were sensitive and insensitive to task environments, which indicated that the *d_4_* - *d_2_* difference was greater in the environment-sensitive group than in the insensitive group (*t*_ACC_(11) = 3.36, *p* = .006; *t*_AI_(11) = 3.15, *p* = .012; fig.4C). There was no evidence of these patterns in any other ROI. Similarly, there was no evidence that the distance between neural representations of the reward options changed between environments when behaviour was not taken into account (see supplementary fig.S5). This suggests that participants represented the options relative to one another in a manner that was linked to their ability to identify the rational task strategy.

Having shown that multivariate patterns of activity in dACC and AI captured critical aspects of the task, we conducted univariate timecourse analysis in these regions to relate them to subcortical ROIs. In contrast to the clear evidence of multivariate pattern change, there was no evidence of univariate activity changes in dACC or AI as a function of experimentally manipulated environment-type (*t_dACC; environment-type_*(26)*=* 1.52, *p*=.282; *t_AI; environment-type_*(26)*=* 0.48, *p*=.636). There was also, similarly, very limited evidence that dACC or AI represented environment-driven policy-switches in the manner of DRN (*t_dACC; pursue-change_*(26)*=*-1.81, *p*=.081; *t_dACC; reject- change_*(26)*= -*2.14, *p*=.085; *t_AI; pursue-change_*(26)*=*0.96, *p*=.346; *t_AI; reject-change_*(26)*=-*2.13, *p*=.085; fig.4D). In combination with earlier analyses of DRN activity, we therefore reasoned that DRN and dACC/AI might constitute a circuit in which cortical components represented options in relation to one another , while DRN implemented the changes in behavioural policy for the options consequent on changes in context. We tested this hypothesis with a series of psychophysiological interaction (PPI) analyses quantifying changes in DRN–dACC/AI connectivity that might reflect communication between regions that varied as a function of the environment. Consistent with this hypothesis, there was a PPI between dACC and DRN whereby coactivation between DRN and dACC was stronger in poor environments, in which environment-driven behavioural changes were most prominent (*t*_PPI; dACC-DRN_(23) = -2.45, *p* = .021; fig.5B). There was no evidence of the same pattern between DRN and AI (*t*_PPI; AI-DRN_(23) = - 0.86, *p* = .400).

**Figure 5.**
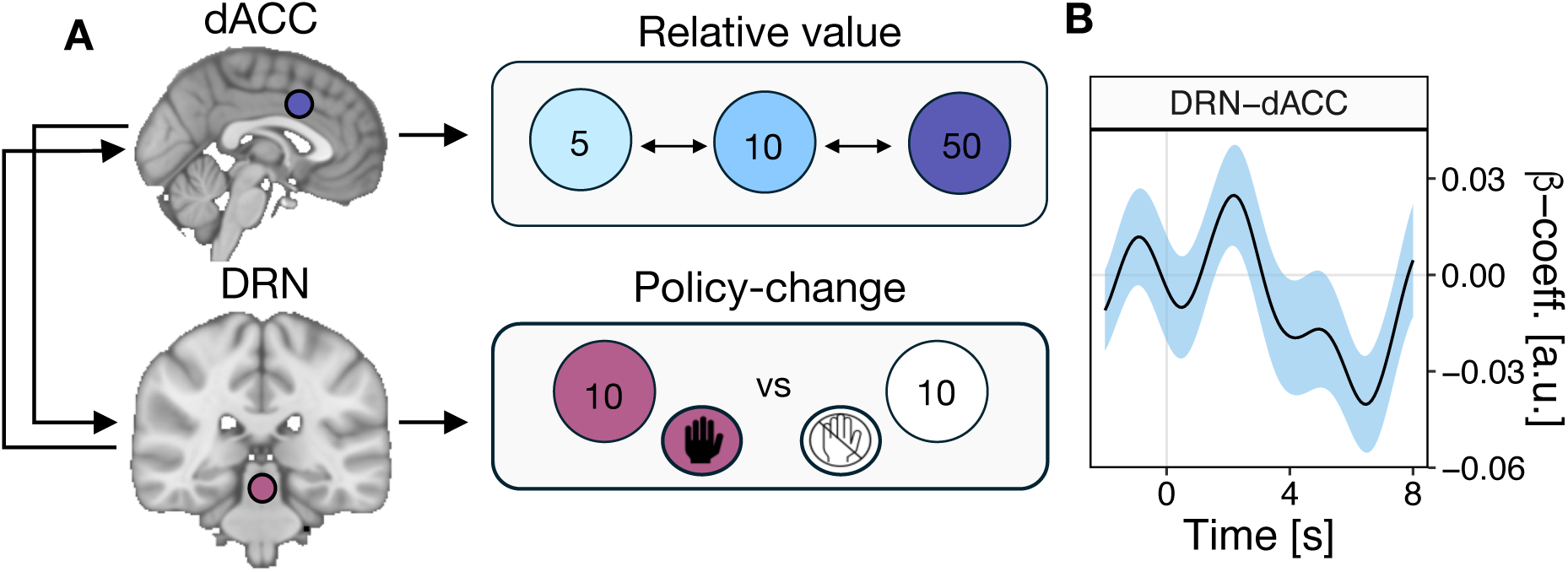
A cortico-subcortical circuit for reconciling behavioural policies with the environment. **(A)** We demonstrate complementary patterns of brain activity in the cortex and dorsal raphe nucleus (DRN) during reward-guided decision-making. In tandem with previous studies, we propose a circuit in which: (i) regions of cortex such as dACC and AI compute the relative value of reward-options given the distribution of recent opportunities and outcomes, and; (ii) DRN implements and/or facilitates changes in behavioural policy that arise from relative value computations. **(B)** Consistent with this hypothesis, we observed a PPI between dACC and DRN as a function of environment-type. The sign of the PPI indicates that there was stronger coactivation between DRN and dACC in poor environments, where behavioural changes were most prominent.

## Discussion

Normative theories of reward-guided behaviour suggest that reward opportunities should be evaluated relative to the context they occur in (Stephens & Krebs, 1986). Here, we show that human behaviour obeys this principle. We gave participants a novel behavioural task involving repeated encounters with the same set of reward options and found that participants were *ceteris paribus* more likely to pursue opportunities in poor environments with few high-value alternatives. fMRI recordings of brain activity linked successful task performance to a distributed neural circuit comprising DRN in the brainstem, and dACC and AI in the cerebral cortex. Within this group, activity in DRN, and no other subcortical nuclei investigated, signalled changes in option-specific behavioural policies, but only insofar as those switches were appropriate to the richness of the environment. By contrast, activity elsewhere,in the neuromodulator system, in MS (Fig.S6), was linked to exploratory switches. Within the DRN-dACC-AI circuit, it was in the dACC and AI that multivariate activity reflected options and their relative similarity in behavioural policy given the environment. In other words, these regions represented roptions in a context-dependent manner. Co-activation between DRN and cortical regions like dACC changed as a function of the environment, and was particularly strong during poor environments where changes in behaviour and brain activity were most prominent. We therefore suggest that DRN, dACC and AI constitute a circuit for reconciling behaviour with the richness of the environment – a circuit in which cortical components maintain representations of options given the current context, and DRN implements changes in behavioural policy relating to these option.

The present work adds to a growing corpus of experiments inspired by behavioural ecology (Constantino & Daw, 2015; Khalighinejad et al., 2020, 2021; Kolling et al., 2012). These studies have shown that human participants are keenly sensitive to background reward-rates during foraging-style scenarios, where their behaviour approximates the predictions of optimal foraging theory. Here, we provide further evidence to this effect by showing that participants pursued a moderately- valuable reward option – the 10-point option – more frequently in poor environments compared to rich environments. A novel feature of our task design was that participants repeatedly encountered the same reward options many times over, and that changes in the environment were engendered by simply manipulating the frequency of each individual option. This meant that we could explicitly track option-specific behavioural policies and their relationship with the richness of the environment. Policy switches were consistent with the behavioural patterns described above, whereby participants switched toward pursuing the 10-point option in poor environments.

Importantly, pinpointing the trials when policy switches occurred allowed us to uncover changes in DRN activity that signalled these switches. This pattern was specific to DRN, and was specific to switches that were appropriate to the richness of the environment. It was further accompanied by activity representing the richness of the environment itself, which emerged gradually over time in the same way as environment-related patterns of decision-making. The joint observation of these neural signals is consistent with recordings of single-neuron activity implicating DRN in environment-driven changes in behaviour, including changes between patience and impulsivity during intertemporal choice (K. Miyazaki et al., 2011; K. W. Miyazaki et al., 2014), exploitation and exploration during reversal learning (Clarke et al., 2004; Grossman et al., 2022; Lottem et al., 2018; Matias et al., 2017), motivation and demotivation during reward seeking (Priestley et al., 2024). The present results provide the first evidence of this phenomenon in the human brain and indicate that DRN has a conserved function of matching the general policy governing behaviour with the general reward statistics of the environment.

Intriguingly, environment-related modulation of both behaviour and DRN activity was more prominent in poor environments compared to rich environments. Given that DRN is the main source of serotonin to the mammalian forebrain, it is tempting to relate this asymmetry to classic theories implicating serotonin in aversive processing (Daw et al., 2002; Jacobs & Azmitia, 1992; Soubrié, 1986), and the ability to adopt aversion- and/or threat-oriented modes of behaviour (Trier et al., 2023). The present results do not implicate DRN activity in aversion *per se*, however. For example, one of the critical DRN activity patterns observed here signalled *pursuits of* – rather than aversion from –the 10-point option in poor environments. Nevertheless, this pattern is broadly consistent with the notion that DRN facilitates behavioural adaptation to adverse circumstances, because pursuits of the 10-point option were more common in poor environments. It is unclear whether this function arises from serotonin neurons specifically, as many DRN neurons are non-serotonergic and it is not possible to discriminate serotonergic from non-serotonergic populations in fMRI recordings (Fu et al., 2010; Jacobs & Azmitia, 1992). Future experiments in human participants could address this by examining whether pharmacological manipulations of the serotonin system influence patterns of behavioural policy change.

In contrast to DRN’s role in policy switches, dACC and AI activity reflected options and their similarity to one another in the task space given the richness of the environment, This accords with previous evidence that these regions track rewards over multiple timesteps and, in dACC’s case, that it evaluates rewards relative to the counterfactual alternatives that might be pursued in the current context (Fouragnan et al., 2019; Hayden et al., 2011; Khalighinejad et al., 2021; Wittmann et al., 2020). Here, we probed this role by testing whether the representational distance between reward options changed across rich and poor environments. This demonstrated that individual differences in the distance between 10-point and 50-point options paralleled the degree of behavioural adaptation to rich and poor environments, suggesting that successful performance of the task was fundamentally linked to warps in the neural representations of options – warps which brought the 10-point and 50-point options closer together when they evoked the same behavioural policy, and pushed them further apart when behaviour diverged. Strikingly similar patterns have been observed in experiments applying RSA to other cognitive domains like navigation, where neural representations of spatial location appear to warp according to an agent’s current and goal locations (Muhle-Karbe et al., 2023). To our knowledge, the present results are the first evidence of the same warping phenomenon in an abstract task space and suggest that warping is a common neural motif of context-dependent computation.

Notably, dACC and AI were the only ROIs with task-related changes in representational distances, and they were selected as ROIs in the first instance because they showed univariate increases in activity during reward-pursuit decisions. It is important to highlight, therefore, that we measured the distance between multivariate patterns of reward-evoked activity with cosine dissimilarity – a method that obviates the influence of univariate changes on representational distance. This means, for example, that environment-driven changes in the representational distance between 10-point and 50-point options is not reducible to the fact that: (i) both 10-point and 50-point options are pursued poor environments, and therefore evoke similar levels of univariate activity, and (ii) the 50-point option is pursued more than the 10-point option in rich environments, leading to divergence in univariate activity. Moreover, the multivariate activity that we recorded in dACC and AI was time-locked to option presentation rather than to participants’ subsequent responses. Nevertheless, given the presence of pursuit-rejection-related activity in dACC and AI, it may be more appropriate to conceive of the multivariate activity in dACC and AI as not simply reflecting the options per se but option-linked behavioural policies. Such a view is consistent with the interactions between DRN and dACC that occurred during the updating of behavioural policies.

Finally, it is worth briefly noting how the present results might relate to affective disorders. Affective disorders are marked by an inability to adapt behaviour to negative outcomes and have been linked to serotonin dysfunction through the efficacy of serotonin specific reuptake inhibitor (SSRI) treatments (Cools et al., 2008; Dayan & Huys, 2009). Although the relationship between serotonin and affect is poorly understood (Cowen & Browning, 2015), the fact that DRN activity signalled behavioural adaptations to poor environments in the present results is broadly consistent with the hypothesis that such disorders can be linked to serotonergic dysfunction. Furthermore, the present focus on general features of behaviour like behavioural policies might be a useful framework for operationalising critical aspects of affective disorders, such as apathy and anhedonia. This perspective on behaviour has not, to our knowledge, been explored in psychopharmacology or computational psychiatry, and might be a fruitful avenue for future experiments.

## METHODS

### Participants

Twenty-nine participants performed the experiment. All were between 18-40 years of age, reported normal or corrected-to-normal vision, and no current diagnosis or treatment for psychiatric or neurological disorder. Participants received £40 for completing the experiment and could earn an additional payment of up to £10 depending on their performance in the decision-making task. All relevant ethical regulations governing research with human participants were observed, and participants provided written informed consent at the beginning of each session. Ethical approval was given by the Oxford University Central University Research Ethics Committee (CUREC) (Ref-Number: MSD-IDREC-R55856/RE001). Two participants were excluded from behavioural analyses because they did not perform the task correctly. An additional two participants were excluded from fMRI analyses due to excessive head motion which prevented accurate registration (motion outliers > 15% of total fMRI volumes).

### Behavioural task

Before performing the experiment in the MRI scanner, participants received written instructions and completed a short practice version of the task. The experiment proper involved five rounds of the behavioural task, and each round was approximately 9 minutes (supplementary fig.1A). Participants were given a 60s break between rounds, meaning that the total duration was approximately 50 minutes.

The behavioural task involved a series of encounters with sequential reward opportunities (fig.1A). The aim was to earn as many points as possible in the fixed task (45 minutes), and participants were incentivised to do this because their payment was partly determined by their performance. In brief, participants were presented with a single reward option on each trial and needed to decide whether to pursue (i.e. accept) or reject it. Pursuing an option incurred a temporal opportunity cost. Given the time-limited nature of the experiment, it was therefore critical to track the frequency of reward options over time – i.e., to anticipate what kind of reward-option they might get in future trials – when deciding whether the opportunity cost was worth incurring. To make the task intuitive, it was presented as a treasure-hunt scenario in which the participant played the role of a ship’s captain searching for treasure opportunities (i.e. reward options; fig.1A). Upon encountering a treasure opportunity, the captain needed to decide whether it was worth the time and effort to sailing toward it (pursue-decision), or to continue to explore the seas elsewhere (reject- decision).

Reward opportunities were drawn from a set of three, discrete options represented by visual stimuli: (1) a low-value option (5 points; bronze medallion) (2) a moderate value option (10 points; silver medallion), and (3) and a high value option (50 points; gold medallion). The value of options was deterministic, constant over time, and known *a priori* to participants before they performed the experiment. The task, thus, did not involve learning about stimulus-reward relationships, or uncertainty about the payout of an option if/when it was chosen. Although option values were constant, the frequency of each option was systematically manipulated to create differences in the richness of the environment. Environments were operationalised via blocks of trials that were time-limited to 4.5 minutes. In rich blocks, the 50-point option was more frequent than the 10-point and 5-point options (fig.1B; *P(Low|Rich)* = 0.16, *P(Middle|Rich)* = 0.33, *P(High|Rich)* = 0.50). In poor blocks, the 5-point option was more frequent than the 10-point and 50-point counterparts (fig.1B; *P(Low|Poor)* = 0.50, *P(Middle|Poor)* = 0.33, *P(High|Poor)* = 0.16].

There were no cues indicating the environment during the task, and participants were not informed that the task was divided into discrete rich and poor environments – all they were told was that the frequency of the options might vary over time. As such, participants needed to learn the richness of the environment by tracking the frequency with which specific options occurred.

Per above, the experiment consisted of five, 9-minute rounds and each round consisted of two 4.5-minute blocks (supplementary fig.S1A). The amount of time remaining in each round was signalled using a timer in the top right-hand side of the screen. Each round featured one rich block and one poor block, and the order of rich/poor blocks within each round was random (supplementary fig.S1A). The precise number of trials completed in each block – and therefore each round – varied depending on how many options were pursued during the allotted time (supplementary fig.S1B; μ(trials)_rich_ = 15.95 ± 1.06, μ(trials)_poor_ = 18.06 ± 1.27, *t*(26)_rich- vs-poor_= 15.52 , *p* <.001). Each trial began with an inter-trial-interval (ITI; *ITI ∼TruncExp*(*μ = 4.5, min=3.5, max=5.5*), during which a treasure chest visual stimulus representing an upcoming reward opportunity descended from the top to the centre of the screen. A visual stimulus – a picture of the medallion – replaced the treasure chest when the chest reached the centre of the screen, thereby revealing the reward-value of the opportunity. Participants needed to consider the reward-option for a short duration (*Opportunity ∼ ∼TruncExp*(*μ = 4.5, min=3.5, max=5.5*) before it displaced either left- or right-ward from the centre. The displacement functioned as a go-cue which indicating that the reward-option was eligible for pursuit. Imposing the go-cue delay created a temporal dissociation between brain activity related to pursue-vs-reject decisions, and the motor processes undertaken to execute the decision via buttonpress. Participants could pursue options by making a button press response within 1s of the go-cue. Different responses were required for leftward and rightward pursuits to prevent anticipatory planning of movements during the go-cue delay. To reject a reward-option, participants needed to withhold any button-press response and let the reward-option pass.

Pursuing an opportunity incurred a temporal opportunity cost via action-outcome (*Action-outcome ∼ TruncExp*(*μ = 4.5, min=3.5, max=5.5*) and reward feedback delays (*Rw-feedback ∼ TruncExp*(*μ = 4.5, min=3.5, max=5.5*). The duration of these delays was equivalent to the time taken by one future encounter – in other words, pursuing an opportunity on the current trial meant foregoing a reward opportunity in the future. The magnitude of this cost was explained to participants during the instructions, and visually indicated during the task via a passing treasure- chest stimulus during the action-outcome delay, which represented an opportunity forgone.

The parameters of the task were carefully chosen to ensure that the optimal strategy was to always pursue the 10-point option in rich blocks, and to never pursue the 10-point option in the poor blocks. Per above, blocks were not signalled to participants, and we therefore did not expect them to display the optimal pattern of behaviour immediately. Instead, we expected that they would move towards the optimal pattern after accumulating experience of each block-based environment over time. Designing the task in this way ensured that there was a reward-maximising rationale for participants to change their behavioural policy toward the 10-point option.

The task was implemented in MATLAB R2022b (Mathworks Inc.) using the Psychophysics Toolbox extension (Kleiner et al., 2007).

### Behavioural analysis

Our initial analysis of behaviour quantified the factors influencing the pursue- vs-reject decision made on each trial. We accomplished this with a series of mixed-effect binomial generalised linear models (GLMs) with participant-identity as a random variable. All GLMs therefore accounted for inter-subject variability in effects of interest.

We first investigated whether pursue-vs-reject decisions were influenced by the richness of the environment, where richness of the environment was operationalised as the experimentally controlled environment-type (rich-vs-poor block) on a given trial:

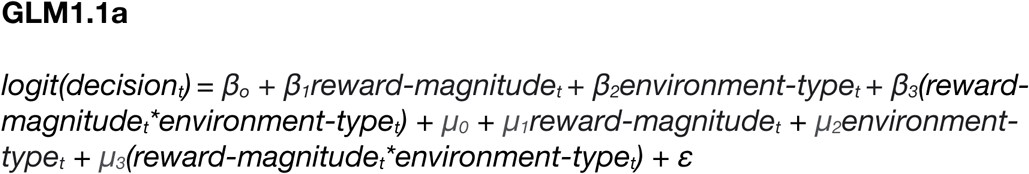

Where β_0-3_ are fixed-effects and μ_0-3_ are random-effects within each subject. We followed this with a GLM that tested the effect of the environment separately for each reward option:

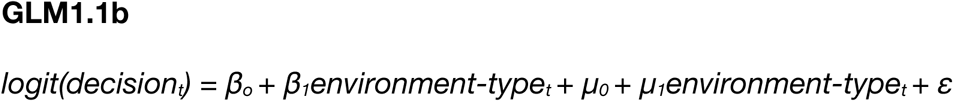

Where β_0-1_ are fixed-effects, μ_0-1_ are random-effects within each participant. Per above, GLM1.1b was fit separately to subsets of trials featuring the 5-point, 10- point and 50-point options, respectively.

We then investigated whether the effect of environment-type on pursue-vs- reject decisions was modulated by the amount of time elapsed within a block (*trial-within-block*). This interaction is consistent with the hypothesis that participants incrementally learned the richness of the environment with experience:

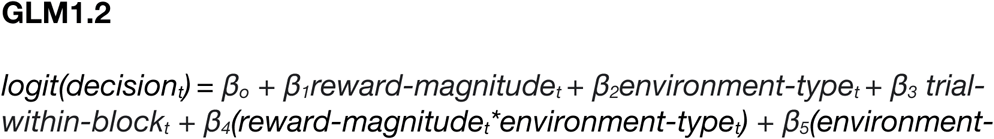

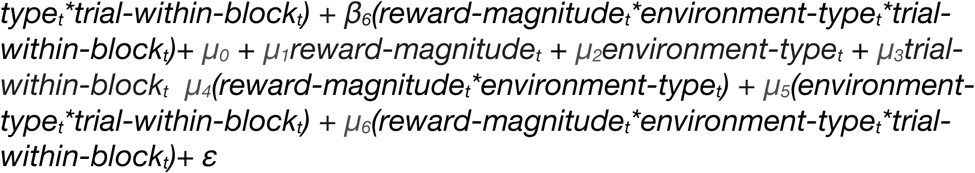

Where β_0-6_ are fixed-effects and μ_0-6_ are random-effects within each subject.

We then tested the same hypothesis from a complementary perspective by operationalising the richness of the environment as the average-value of the reward- options encountered on the previous five-trials (*average-value*):

#### Average-value

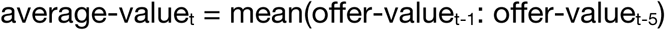

Operationalising the richness of the environment in this way captured a subject’s experience of the task, instead of relying on the task’s underlying generative structure. We investigated the effect of average-value on pursue-reject decisions with the following GLM:

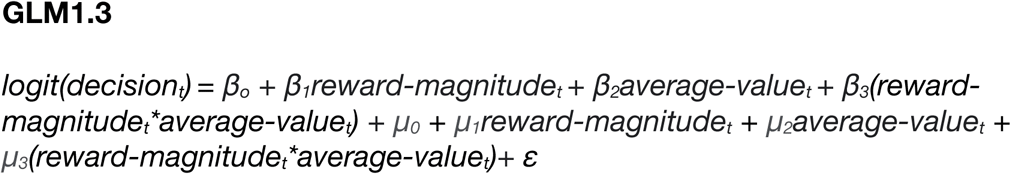

Where β_0-3_ are fixed-effects and μ_0-3_ are random-effects within each subject.

Because the task featured three discrete reward-options that occurred many times over, we were able to precisely identify changes in a subject’s behavioural policy toward each option (where behavioural policy indicates an option-specific pursue-vs-reject strategy; Fig.2A). We first tested the hypothesis that participants formed option-specific policies by examining whether pursue-vs-reject decisions were autocorrelated over consecutive encounters with a particular reward-option:

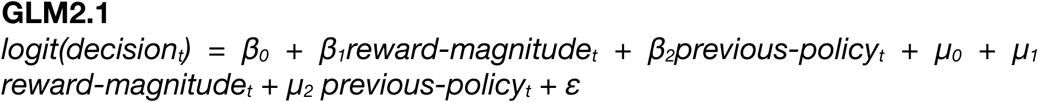

Where β_0-2_ are fixed-effects and μ_0-2_ are random-effects within each subject. Here, previous-policy indicates the pursue-vs-reject decision the last time that the reward option on trial *t* was encountered. We confirmed that behavioural policies were option-specific by testing the effect of the pursue-vs-reject decision taken on the preceding trial (i.e. *t*-1) regardless of which reward option occurred:

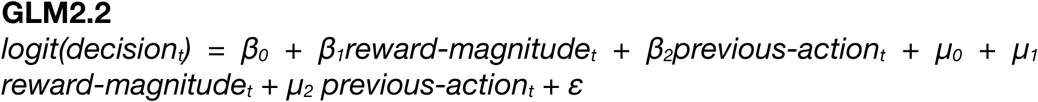

Where β_0-2_ are fixed-effects and μ_0-2_ are random-effects within each subject. Together, GLM2.1 and GLM2.2 allowed us to test whether temporal structure in behaviour was option-specific, or whether participants also had autocorrelated predispositions to pursue-vs-reject options in general, regardless of reward-option identity.

We then operationalised policy-switches as trials on which a subject changed their pursue-vs-reject decision across successive encounters with a particular option – for example, if a subject rejected the 10-point option on trial *t=1* and subsequently accepted the 10-point option on trial *t=5*, trial *t=*5 was coded as policy-switch event. We also defined trials along an action dimension, where an action-switch occurred when participants changed their pursue-vs-reject decision across consecutive trials (regardless of which reward-option appeared). This allowed us to compare option-specific tendencies (i.e. policy dimension) in purse/reject decisions with general tendencies (i.e. action dimension) in pursue-vs-reject decisions.

After defining each trial along policy-switch and action-switch dimensions, we conducted a series of analyses to identify the factors predicting policy-switches. Because the policy-switch status of each trial was binary (switch=1, stay =0), we used mixed-effect binomial generalised linear models (GLMs) where the dependent variable was the policy-switch status of each trial, and participant-identity was a random variable.

We first investigated whether the probability of policy-switches differed as a function of reward-option:

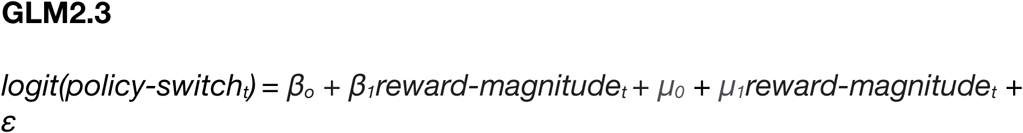

Where β_0-1_ are fixed-effects and μ_0-1_ are random-effects within each subject. We then investigated whether the direction of policy-switches for the 10-point option was modulated by environment-type. To do this, we separated policy- switches into two different types: (1) *reject-switches*, when subjects rejected the 10- point reward-option but pursued it on the preceding encounter, and (2) *pursue- switches,* where subjects pursued the 10-point option but rejected it on the previous encounter. The optimal strategy for performing the task involved pursuing the 10- point option in poor environments and rejecting the 10-point option in rich environments. Insofar as behaviour approximated this strategy, we expected that pursue-switches would occur more frequently in poor environments and reject- switches would occur more frequently in rich environments. We tested these hypotheses with the following GLMs:

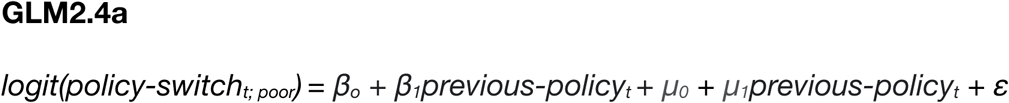

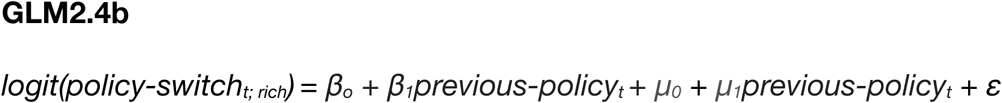

Where β_0-1_ are fixed-effects and μ_0-1_ are random-effects within each participant. GLM2.4a and GLM2.4b were fit to trials featuring the 10-point option. Note that although GLM2.4a and GLM2.4b have the same formula, GLM2.4a was fit only to trials from poor blocks and GLM2.4b was fit only to trials from rich blocks. We then tested the same hypothesis in a complementary way with the following GLMs:

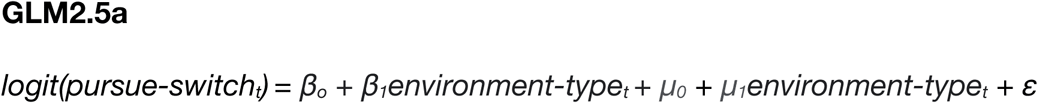

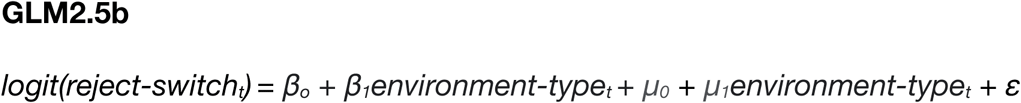

Where β_0-1_ are fixed-effects and μ_0-1_ are random-effects within each participant. GLM2.5a and GLM2.5b were fit to trials featuring the 10-point option. We then tested the same hypotheses in an analysis where the richness of the environment was operationalised as the average-value of the reward-options encountered on the previous five-trials:

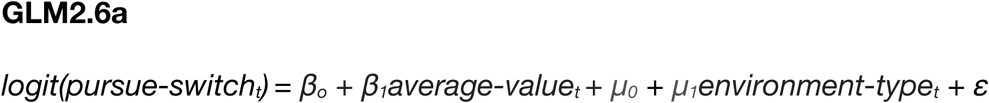

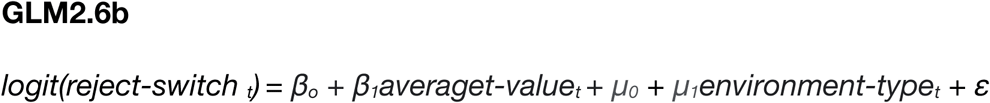

Where β_0-1_ are fixed-effects and μ_0-1_ are random-effects within each participant. GLM2.6a and GLM2.6b were fit separately to subsets of trials featuring the 5-point, 10-point and 50-point options respectively. All behavioural analyses were performed in *R* (R Core Team, 2023) using the *lme4* implementation of mixed-effects GLMs (Bates et al., 2015).

### Imaging data acquisition

Structural and functional MRI data was collected with a Siemens 7 Tesla MRI scanner. High-resolution functional data were acquired with a multiband gradient-echo T2* echo planar imaging sequence with 1.5mm isotropic voxels, multiband acceleration factor 2, repetition time (TR) = 1.775s, echo time (TE) = 17.8ms, flip angle = 66°, and GRAPPA acceleration factor 2. The parameters were selected to maximise signal-to-noise ratio in subcortical areas. To accommodate the high temporal and spatial resolution of the protocol, functional scans had a limited field of view (FOV) oriented at 30 degrees with respect to the AC-PC line (66 slices with a coverage of 99 mm). The FOV captured all regions of interest in the midbrain, brainstem and cortex. Before acquiring the task-related functional scan, we acquired a pre-saturation single-measurement, whole-brain functional scan with the same orientation. The pre-saturation scan was used to facilitate registration of the limited- FOV task-related functional scan to the whole brain.

Structural data were acquired using a T1-wieghted MP-RAGE sequence with 0.7mm isotropic voxels, GRAPPA acceleration factor 2, TR = 2200ms, TE = 3.02ms, and; inversion time (TI) = 1050ms. To correct distortions arising from inhomogeneities in the magnetic field, a fieldmap sequence was acquired with 2mm isotropic voxels, TR = 620ms, TE1 = 4.08ms, and TE2 = 5.1ms. To account for the effects of physiological noise on functional MRI data, participants were fitted with a pulse oximeter and respiratory bellows that acquired cardiac and respiratory timeseries at 50Hz using a BioPac MP160 device (BIOPAC Systems Inc., USA).

### fMRI data preprocessing

Preprocessing of fMRI data was performed with the FMRIB Software Library (Jenkinson et al., 2012; Smith et al., 2004). The Brain Extraction Tool (Smith, 2002) was used to separate brain from non-brain matter in structural and functional images. Functional images were normalised, spatially smoothed (Gaussian kernel with a 3mm full-width half-maximum) and temporally high-pass filtered (3 dB cut-off = 100s), and artefacts arising from head motion were removed using MCFLIRT (Jenkinson et al., 2002). Registration of task-related functional images to Montreal Neurological Institute (MNI)-space was performed in three stages:

1. The task-related limited-FOV EPI was registered to the pre-saturation whole-brain EPI using FMRIB’s Linear Image Registration Tool with 6 degrees of freedom transformation.
2. The whole-brain EPI was registered to the subject-specific structural images using Boundary-Based Registration (BBR) incorporating fieldmap correction (Greve & Fischl, 2009).
3. Subject-specific structural images were registered to a 1mm resolution Standard MNI template with FMRIB’s Non-linear Registration Tool (FNIRT; (Jenkinson et al., 2012)).

### Whole-brain fMRI data analysis

Statistical analysis of whole-brain functional data was performed at two levels using FMRIB’s Expert Analysis Tool (FEAT; (Jenkinson et al., 2012)). In the first level, a univariate general linear model was used to compute parameter estimates for each regressor in each participant (Woolrich et al., 2001). Contrast and variance estimates for each parameter in each participant were subsequently combined in a mixed-effects analysis conducted at the second level (FLAME 1 + 2), where subject-identity was a random effect (Woolrich et al., 2004). Significance testing was performed with cluster-correction, a cluster significance threshold of *p* = .001, and a voxel inclusion threshold of *z* = 3.1. Data were pre-whitened before analysis to account for temporal autocorrelations in BOLD signal.

We performed two-whole brain analyses. The first GLM identified voxels where BOLD signal represented the pursue-vs-reject decision made on each trial. Note that the pursue-vs-reject decision made on each trial is regressed at two different timepoints: (1) at the time of opportunity onset, and (2) at go-cue onset. These events were separated in time by a jittered interval drawn from *Opportunity ∼ TruncExp*(*μ = 4.5, min=3.5, max=5.5)*, meaning that neural activity at each time was temporally decorrelated. The second GLM identified voxels where BOLD signal represented the reward-magnitude of the reward-option on each trial. We could not include both pursue-vs-reject and reward-magnitude regressors in the same GLM because they were highly correlated (*r* = 0.64). The formulas for GLMs were:

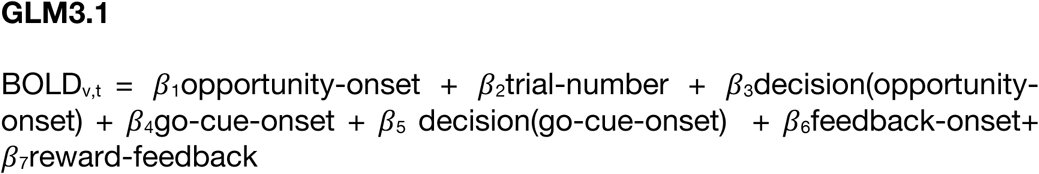

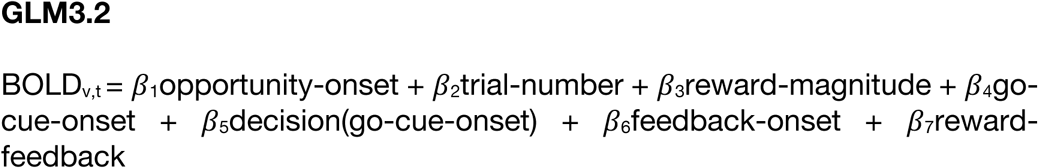

Where BOLD_v,t_ is a *t* x *v* matrix containing the BOLD signal measurements at time *t* in voxel *v*. The regressors are as follows: *opportunity-onset* is an unmodulated regressor representing the occurrence of the reward-option stimulus on each trial; *trial-number* is a parametric regressor representing the number of trials completed in the experiment as a proxy for the passage of time, time-locked to the opportunity presentation; *decision(opportunity-onset)* is a binary regressor reflecting the pursue- vs-reject decision on each trial, timelocked to the onset of the reward-option stimulus; *reward-magnitude* is a parametric regressor reflecting the value ∈ {5, 10, 50} of the reward-option timelocked to opportunity-onset; *go-cue-onset* is an unmodulated regressor representing the occurrence of the go-cue stimulus on each trial; *decision(go-cue-onset)* is a binary regressor reflecting the pursue-vs-reject decision on each trial, timelocked to the onset of the go-cue stimulus; *feedback-onset* is an unmodulated regressor representing the occurrence of reward-feedback received on trials when the opportunity was accepted; *reward-feedback* is a parametric regressor with three levels (5, 10, 50) representing the reward-option, and time-locked to feedback presentation on trials where opportunities were accepted.

All regressors were boxcar functions with a constant duration of 0.1s and convolved with a double-gamma hemodynamic response function. Further non-task confound regressors were added to reduce noise in BOLD signal, including: (1) head motion parameters estimated using MCFLIRT during pre-processing (Jenkinson et al., 2002); (2) regressors for voxel-wise estimates of physiological noise arising from cardiac and respiratory activity, estimated using FSL’s Physiological Noise Monitoring (PNM) tool (Brooks et al., 2008), and; (3) regressors for motion outliers, indicating volumes with head motion that could not be corrected with linear methods.

### ROI time course fMRI analysis

Additional analysis of brain activity was conducted on epoched time-course activity extracted from regions of interest (ROIs). Anatomical ROIs were constructed for a series of brainstem and midbrain nuclei previously implicated in reward-guided decision-making including dorsal raphe nucleus (DRN), Locus Coeruleus (LC), Medial Septum (MS), a combined mask covering the Substantia Nigra and Ventral Tegmental Area (Midbrain Dopaminergic Nuclei; MDB), and Habenula (Hb). The LC mask derived from a brain atlas developed by Pauli and colleagues (Pauli et al., 2018). DRN, MDB, Hb and NB masks were manually drawn in MNI-space based on an atlas of the human brain atlas (Mai et al., 2016). The DRN mask was drawn in consultation with the anatomical guidelines of Kranz and colleagues (Kranz et al., 2012) to ensure specificity to the dorsal portion of raphe nucleus. Standard-space masks were transformed to subject-specific structural space by applying a standard-to-structural warp field. To ensure conformation between anatomical masks and subject-specific neuroanatomy, subject-specific masks were manually checked and edited by two experimenters and evaluated for inter-rater reliability by a third experimenter. Structural-space masks were then transformed from structural to functional space by applying a structural-to-functional affine matrix, and were subsequently thresholded, binarized and dilated by one voxel.

To prepare ROI data for analysis, the filtered time-series of BOLD signal from each voxel was averaged, normalised and up-sampled by a factor of 20 with spline interpolation (Behrens et al., 2008). Up-sampled timeseries were then epoched in 10s windows starting 2s before and ending 8s after reward-option onset on each trial. Timeseries data was analysed by fitting a GLM with ordinary least squares (OLS) at each timepoint in each epoch. The statistical significance of parameters in GLMs was assessed using a leave-one-out procedure that avoided temporal biases in the selection of peak-effects (Kolling et al., 2018). The procedure involved estimating, for each regressor *r* in each subject *s*, the time *t* at which the mean effect of *r* was greatest in the remaining *N*-1 participants. The effect of *r* in *s* was then estimated by calculating the effect size for *r* at time *t* in subject *s*. This was repeated iteratively for each subject, leaving an *N*x1 vector of parameter estimates that were not biased by the timecourse of data of individual participants. We assessed statistical significance of subject-specific parameter estimates by running a single-sample t-test on the vector of parameter estimates. All t-tests were two-sided. Whenever analyses involved three or more ROIs, we maintained the family-wise error rates at α=.05 using the Holm-Bonferroni procedure for multiple comparisons(Holm, 1979).

We tested which ROIs represented policy-switches with the following GLM:

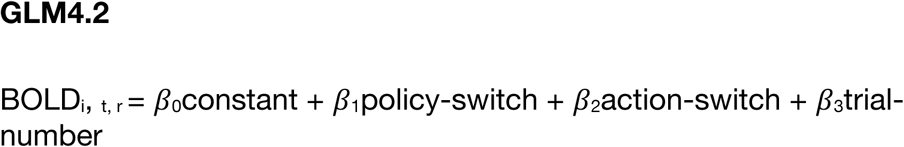

Where *policy-switch* is a binary variable indicating whether a policy-switch occurred on trial *t*, and *action-switch* is a binary variable indicating whether an action-switch occurred on trial *t* (see fig.2A for definitions).

To investigate brain activity related specifically to task-driven policy-switches in behaviour, we classified policy-switch events as either *congruent* or *incongruent* in relation to the environment they occurred in. Recall that the optimal strategy for preforming the task was to pursue the 10-point opportunity in poor environments and reject it in rich environments. Accordingly, we coded pursue-switches in poor environments as congruent and pursue-switches in rich environments as incongruent. Correspondingly, reject-switches in rich environments were congruent and reject-switches in poor environments were incongruent.

We used these four event types as regressors in the following series of GLMs:

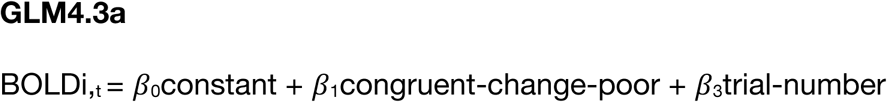

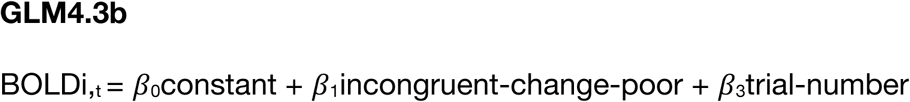

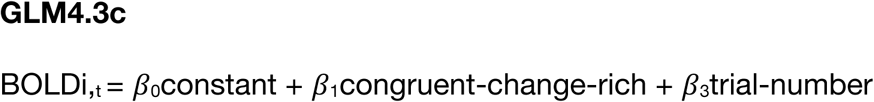

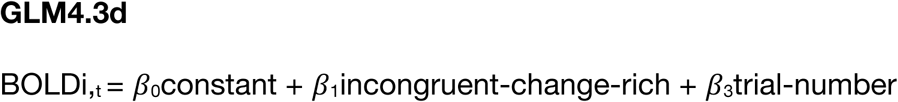

Where *congruent-change-poor* is a binary variable indicating whether a congruent policy-switch in a poor environment occurred on trial *t*, *incongruent-change-poor* is a binary variable indicating whether an incongruent policy-switch in a poor environment occurred on trial *t*, *congruent-change-rich* is a binary variable indicating whether a congruent policy-switch in a rich environment occurred on trial *t*, and *incongruent-change-rich* is a binary variable indicating whether an incongruent policy-switch in a rich environment occurred on trial *t*. We tested differences in neural activity related to these event-types by performing paired-sample t-tests on the effect-sizes for different event types.

Per above, the optimal strategy in the task involved changing behavioural- policy toward the 10-point option depending on the environment. Conversely, there was no task-driven reason for changing behavioural-policy toward the 5-point option, which should always have been rejected. To further test for brain activity related to task-driven policy-switches, we separately analysed the effect of policy-switches on trials featuring the 10-point and 5-point options. We did this by performing the analysis in GLM4.3a and GLM4.3c for subsamples of data containing 10-point option trials and the 5-point option trials, respectively.

### Representational similarity analysis

We further investigated neural representations of the task environment using representational similarity analysis (RSA; (Kriegeskorte et al., 2008)). RSA was performed on the following cortical ROIs: anterior insular cortex (AI), dorsal anterior cingulate cortex (dACC), lateral frontopolar cortex, subgenual anterior cingulate cortex, (sgACC) and pregenual anterior cingulate cortex (pgACC) and dorsomedial prefrontal cortex (dmPFC). AI, dACC and lFPC were selected because they exhibited clusters of activity related to the pursue-vs-reject contrast in whole-brain fMRI analysis (GLM3.1; see supplementary table 1). Similarly, sgACC was selected because it exhibited activity related to the reward-magnitude contrast in a whole-brain fMRI analysis (GLM3.2; see supplementary table 2). dmPFC and pgACC were selected based on functional activations in previous studies (CITATION). Each ROI was constructed as a sphere with 7mm radius centred on the voxel with the highest cluster-corrected β-weight.

We used RSA to compare differences in the neural representation of reward- options between rich and poor environments. We predicted that the distance between option-specific neural representations would mirror changes in the relative-value of reward options given the environment – i.e., the 10-point would be closer to the high-option in poor environments compared to rich environments, and closer to the low-option representations in rich environments compared to poor environments (fig.4A). The analysis consisted of two stages: (1) quantifying the multi-voxel patterns of activity evoked by 5-point, 10-point and 50-point reward options in rich and poor blocks of the experiment, and; (2) computing the distance between multi-voxel representations, and quantifying how distances changed as a function of environment-type.

We measured multi-voxel representations of reward-options using a GLM implemented in FSL FEAT. fMRI data pre-processing followed the pipeline described in *fMRI data preprocessing* above except that spatial smoothing was omitted to maximise voxel-specific information. Omitting spatial smoothing prevented accurate registration for two participants, who were therefore excluded from the analysis (*N_RSA_* = 25). The GLM quantified the multi-voxel patterns of activity evoked by each reward- option with separate binary predictors indicating the onset of each option in each block (i.e., 30 predictors in total). It had the following formula:

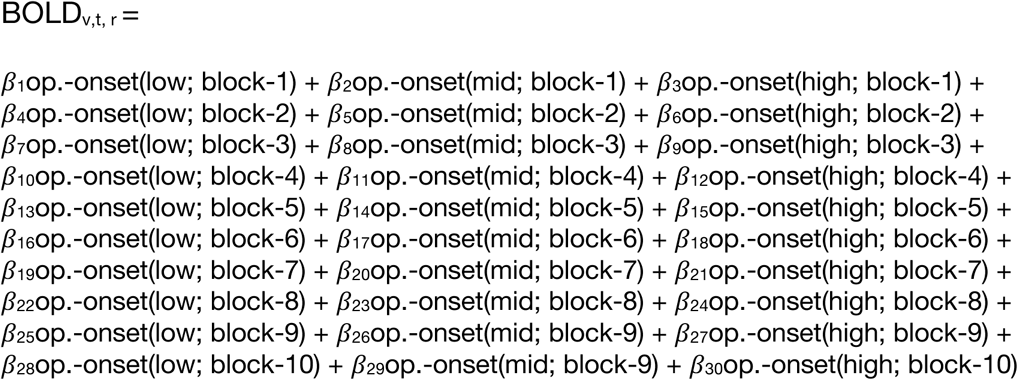

BOLD_v,t,r_ is a *t* x *v* x *r* matrix containing the BOLD signal measurements at time *t* in voxel *v* in ROI *r*. β_1-30_ are unmodulated regressors representing the occurrence of a given reward-option in a given block (e.g. option-onset(low; block-1) indicates onsets of low-option in the first block of the experiment). All option-onset regressors were implemented as boxcar functions with a constant duration of 0.1s and convolved with a double-gamma hemodynamic response function. Confound regressors for physiological noise and motion artefacts were included, as per whole-brain fMRI analysis.

For each ROI *r*, and each predictor *i* in β_1-30_, GLM5.1 produced a vector of *v* regression weights captured the effect of β_i_ at each voxel *v* in *r*. Each vector of regression-weights constituted a multivariate neural representation of a reward- option. We then quantified pairwise distances between multivariate representations using cosine dissimilarity – i.e., the complement of the cosine-angle between vectors. We used cosine dissimilarity instead of standard parametric (e.g. Pearson’s R) and non-parametric (e.g. Kendall’s Tau) methods to avoid artefacts arising from univariate differences in activity (see Discussion for further details). Our analysis focussed on four representational distances; (i) *d_1_*, the distance between 5-point and 10-point options in poor environments; (ii) *d_2_*, the distance between 10-point and 50-point options in poor environments; (iii) *d_3_*, the distance between 5-point and 10-point options in rich environments, and; (iv) *d_4_*, the distance between 10-point and 50-point options in rich environments.

To avoid biases arising from the temporal autocorrelation of BOLD signal, distance calculations were performed by comparing option-specific representations between non-contiguous blocks of the same environment-type (see supplementary sig.S5). For example, to calculate *d_1_* – the distance between 5-point and 10-point options in poor environments – we might compare 𝛽_11_op.-onset(mid; block-4) vs 𝛽_16_opportunity-onset(low; block-6), where blocks 4 and 6 are poor environments that are 1-block apart (block-space, hence). Because the analysis was performed to compare changes in representational distance between environments, we excluded representational distance scores if their block-space was not present for both rich and poor environments. To avoid overrepresentation of specific block-space values in representational distance scores, we calculated the mean representational distance at each value of block-space for each environment. For final representational distances, we calculated the mean representational distance over all levels of block- space in each environment. This process was performed separately for each ROI in each participant.

Once subject-specific *d_1_*, *d_2_*, *d_3_*, and *d_4_*, scores were calculated, we tested our hypotheses by taking the difference between environment-specific distance scores (e.g. H1_10point-vs-5point_ = d_1_-d_3_; H2_10point-vs-50point_ = d_4_ – d_2_). We tested the relationship between representational distances and behaviour by performing Pearson’s *R* correlations between H1 and H2 scores and the degree of behavioural change for the 10-point option between environments [Δ(pursue-rate = pursue-rate_poor_ – pursue- rate_rich_]. We supplemented this by dividing participants into two groups based on their degree of behavioural change: (i) a sensitive group, where (pursue-rate_poor_ – pursue- rate_rich_ ≥ 0), and; (ii) an insensitive group, where (pursue-rate_poor_ – pursue-rate_rich_ < 0). We then tested whether H2 scores were different between groups using a two-sample t-test. In addition, we investigated differences in representational distance regardless of behaviour by testing H1 and H2 against 0 using single-sample t-tests. These analyses were performed separately for each ROI. The family-wise error rate was controlled separately for t-test and correlation analyses using the Bonferroni-Holm correction for multiple comparisons (Holm, 1979).

---

**Figure S1.**
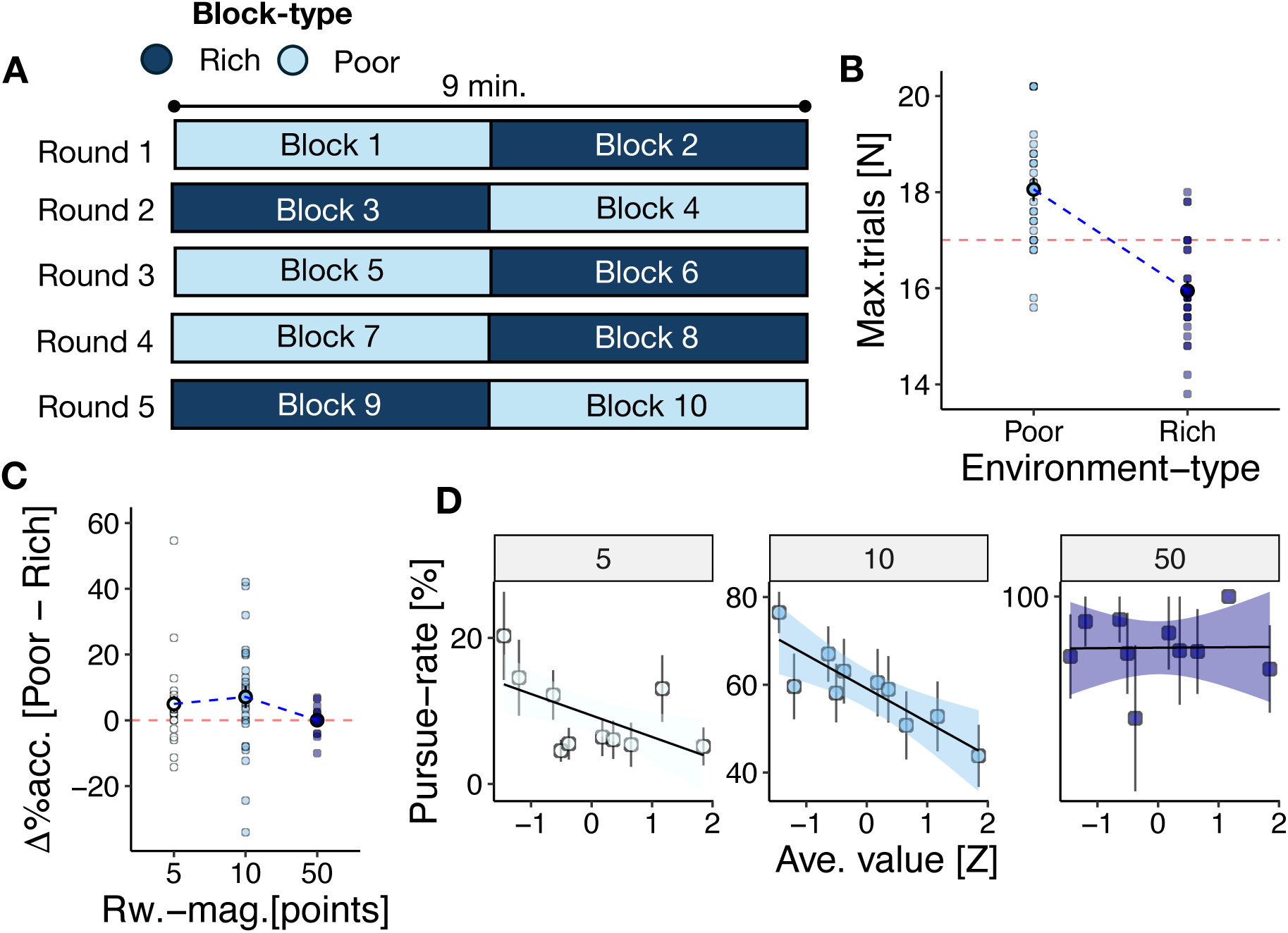
Additional details of the task and behaviour. **(A)** Diagram of the experiment’s round and block structure in an example session. Each experimental session involved five, 9 minute rounds of the behavioural task. Each round of the behavioural task involved two time-limited 4.5 minute blocks. Each round included one rich block and one poor block, which were randomised in order. **(B)** Because blocks were time-limited, and the duration of each trial depended on whether an opportunity was pursued or rejected, the number of trials in each block was variable. Overall, participants performed more trials in poor blocks compared to rich blocks because they pursued fewer opportunities in poor blocks (*t*_poor-vs-rich_(26) = 15.52, *p* <.001). Dots show participant-level mean number of trials in poor and rich blocks **(C)** A GLM predicting the probability of reward-pursuit indicated main effects of reward-magnitude (β_reward-magnitude_= 6.12, SE = 0.25, *p* < .001) and environment-type (β_environment-type_= -2.03, SE = 0.13, *p* < .001), in addition to a two-way interaction between these predictors (β_reward-magnitude*environment- type_= -3.09, SE = 0.19 *p* < .001). Follow-up tests on the interaction indicated that the effect of environment-type was contingent on reward-magnitude: as per the main manuscript, participants were more likely to pursue 10-point (β_environmen-type_ = -0.19, SE = 0.09, *p* = .045) opportunities in poor environments relative to rich environments, but there was no evidence of environment-type changes for the 5-point (β_environmen-type_ = -0.58, SE = 0.31, *p* = .062) or 50-point (β_environmen-type_ = -0.07, SE = 0.24, *p* = .767) opportunities as a function of environment-type. Dots indicate participant-level difference-scores [Δ = μ(pursue)_poor_ – μ(pursue)_rich_] at each level of reward-magnitude. **(D)** Similarly, predicting the probability of reward-pursuit as a function of the average-value of recent opportunities (where ave.- value_t_ = mean(reward-magnitude_t-1_: mean(reward-magnitude_t-5_); see *Methods*) indicated that participants were more likely to pursue 10-point (β_average-value_ = -0.33, SE = 0.06, *p* = .003) and 5-point (β_average-value_ = -0.40, SE = 0.18, *p* = .021) opportunities as average-value decreased. There was no evidence of this effect for 50-point opportunities (β_average-value_ = -0.14, SE = 0.47, *p* = .765). This is consistent with the hypothesis that participants changed their behaviour as a function of recently encountered opportunities. Dots and error bars indicate mean ± SEM pursue-rate in deciles of average-value.

**Figure S2.**
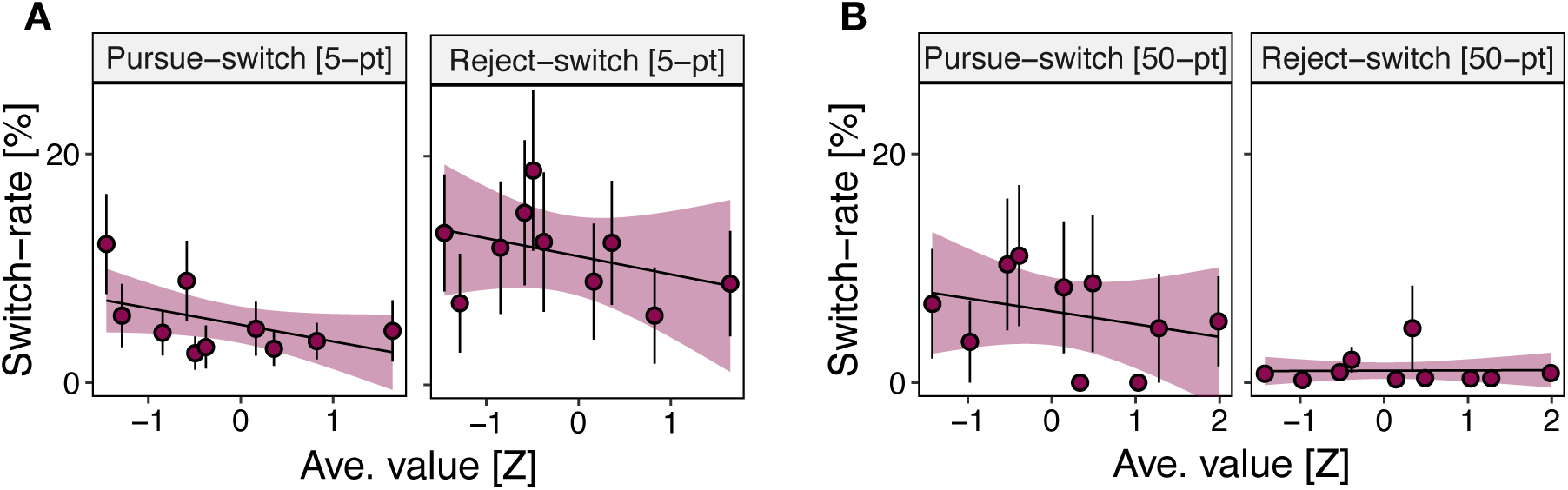
Policy-switches for 5-point and 50-point opportunities are not related to the richness of the environment. Policy-switches for 10-point opportunities were influenced by the richness of the environment. In particular, pursue-switches were more likely to occur as the average-value of recent opportunities decreased (fig.2E). In contrast, there was no evidence that pursue-switches or reject- switches were influenced by average-value for 5-point opportunities (**A**; β_pursue-switch_ = -0.26, SE = 0.14, *p* =.071; β_reject-switch_= -0.03, SE = 0.47, *p* = .873) or 50-point opportunities (**B**; β_pursue-switch_ = -1.36, SE = 0.84, *p* =.107; β_reject-switch_= -0.12, SE = 0.24, *p* = .606). Instead, policy-switches for these opportunities appeared to occur at a constant stochastic rate. In both panels, dots and error bars indicate mean ± SEM switch-rate in deciles of average-value.

**Figure S3.**
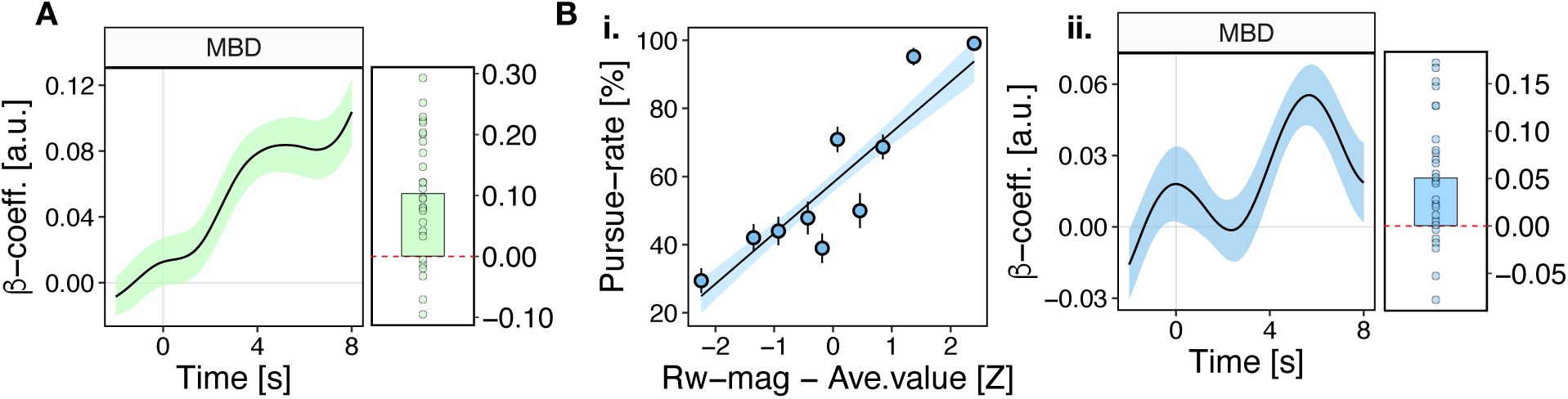
Activity in midbrain dopaminergic nuclei represents pursue-vs-reject decisions, and the value-difference between current and recent reward-opportunities. **(A)** BOLD signal in the MBD ROI was strongly modulated by the pursue-vs-reject decision made on each trial, consistent with the link between dopaminergic nuclei and action initiation (*t_MBD; pursue-vs-reject_* (26)*=*5.21, *p*<.001). **(B)** Although BOLD signal in the MBD ROI did not covary with environment-type or average-value, it was modulated by the value-difference between the reward opportunity on the current trial and the average-value of recent opportunities [i.e. value-difference_t_ = reward-magnitude_t_ – average-value_t_]. This response is reminiscent of a reward-prediction error in the sense that the average-value of recent trials constitutes a simple prediction for the prospective reward-value of future trials *t_MBD; value- difference_*(26)*=*3.96, *p*<.001. Importantly, participants were also more likely to pursue reward-opportunities as a function of value-difference (β_value-difference_= 0.90, SE = 0.07, *p* <.001). In this way, the value-difference signal in MBD is further consistent with the hypothesis that it invigorates reward-guided behaviour. In **(B-i)**, dots and error bars indicate mean ± SEM pursue-rate in deciles of value-difference. In **(A)** and **(B-ii)**, lines and shadings of the time course graph show mean ± SEM of β weights across participants. In effect-size graph, bar show sample-mean magnitude of the peak regression weight in the timecourse according to an unbiased leave-one-out procedure (see *Methods*). Dots indicate peak regression weights for individual participants.

**Figure S4.**
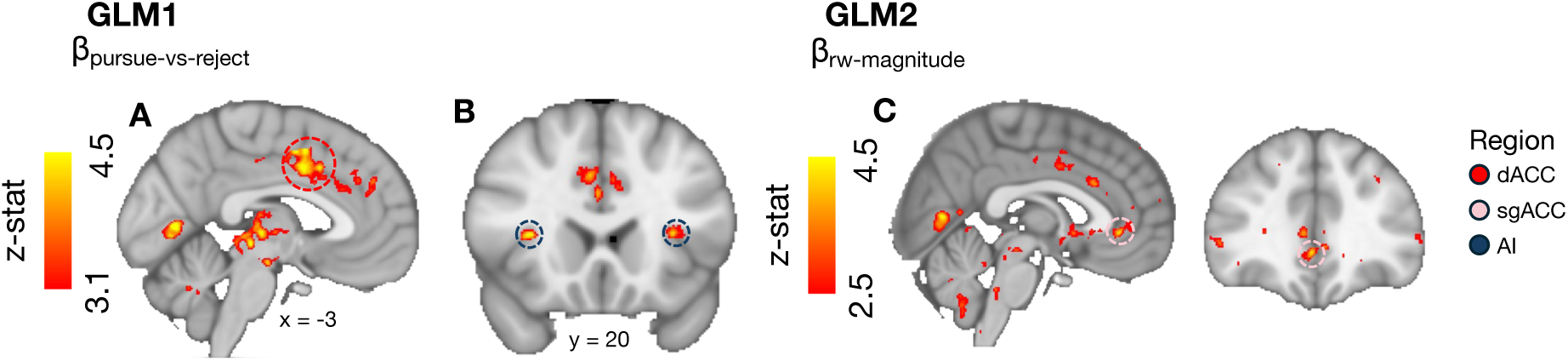
A whole-brain fMRI analysis identifies regions of interest for representational similarity analysis. We conducted representational similarity analysis (RSA) to probe patterns of brain activity corresponding to the value of a reward opportunity relative to the environment. We identified regions of interest (ROIs) for RSA partly using whole-brain fMRI GLMs. In the first whole-brain analysis (GLM3.1), we searched for activity related to the pursue-vs-reject decision made on each trial. This indicated **(A)** a prominent series of clusters in dorsal anterior cingulate cortex (dACC) centred on the cingulate sulcus and extending dorsally into supplementary and presupplementary motor areas (*z*-max = 4.95; MNI-coordinates = [-1, 5, 38]). In addition **(B)**, there were bilateral clusters in anterior insular cortex (AI) at the border with frontal operculum (left z-max = 3.89, MNI-coordinates = [-32, 10, 10]; right z-max = 4.36, MNI-coordinates = [43, 9, 3]). In the second whole-brain analysis (GLM3.2), we searched for activity related to the factors which influenced pursue-vs-reject decisions. This identified **(C)** a cluster of activity in subgenual anterior cingulate cortex (sgACC) related to the reward-magnitude of the opportunity on each trial (*z*-max=3.87, MNI coordinates = [2, 36, -3]). The z-score for the cluster was below the conventional cluster-correction threshold of *z=*3.1. However, we selected it as an ROI because sgACC and neighbouring areas like perigenual anterior cingulate cortex (pgACC) have previously been linked to the neural representation of other aspects of subjective value, which might share similarities with the relative-value concepts that we investigate here.

**Figure S5.**
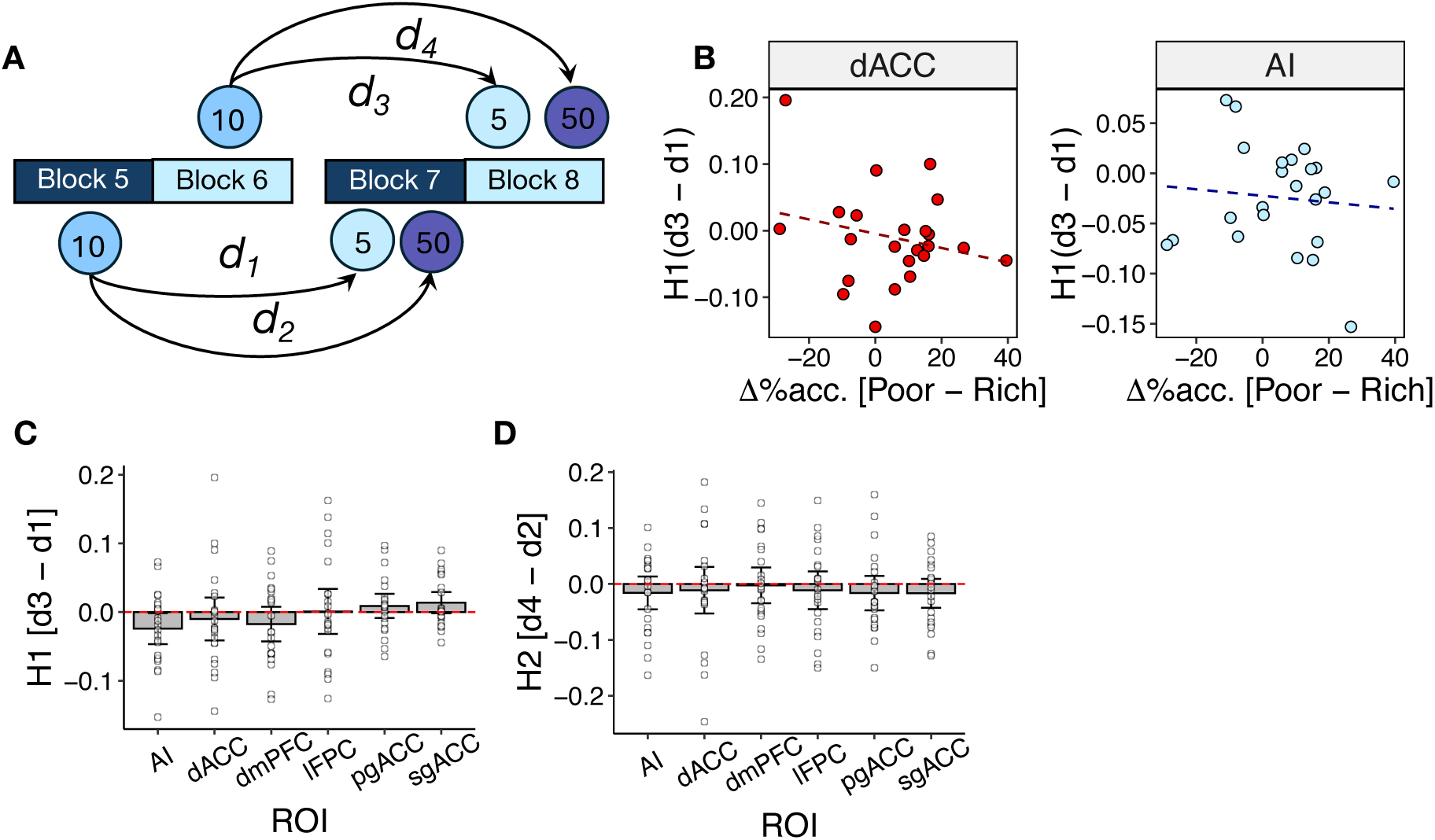
Representational similarity analysis implementation and further results. **(A)** Diagram of strategy for calculating representational distances in representational similarity analysis (RSA). The RSA approach involved comparing the pairwise representational distances separating reward options between rich and poor blocks. To calculate representational distances in the first instance, we compared the similarity of multi-voxel reward-evoked patterns of activity. To avoid similarity artefacts arising from temporal autocorrelations in BOLD signal, we calculated distances using multi-voxel patterns from non-contiguous blocks (see diagram). For example, to estimate the distance between 5-point and 10-point options during poor blocks, we might compare the similarity of 10-point evoked patterns of activity in Block 5 (a poor block) to 5-point evoked patterns of activity in Block 7 (also a poor block). See *Methods* for further details. **(B)** RSA indicated that the H2 – the representational distance between 10-point and 50-point options – in dACC and AI was correlated with rational performance of the behavioural task (fig.4C). In contrast, H1 – the representational distance between 10-point and 5-point options – was not correlated with behaviour in dACC or AI (*t*_dACC_(21) = -1.12, *p*=.271; *t*_AI_(21) = -0.45, *p*=.650), nor in any other ROI. Dots indicate individual participants. Lines represent line-of-best-fit. **(C)** There was no evidence that representations of 10-point and 5-point options were different between poor and rich blocks (i.e. H1 ≈ 0) in any ROI. **(D)** Similarly, there was no evidence that representations of 10-point and 50-point options were different between poor and rich blocks (i.e. H2 ≈ 0). Together with the result in fig.4C, this suggests that the neural representation of reward-options only obeyed changes in the relative value of reward-options in participants who adopted the rational task strategy. In **(C)** and **(D)**, bar and errorbars indicate mean ± 95%CI H1 score in each ROI (n.b. CI not corrected for multiple comparisons). Dots indicate H1 values for individual participants.

**Figure S6.**
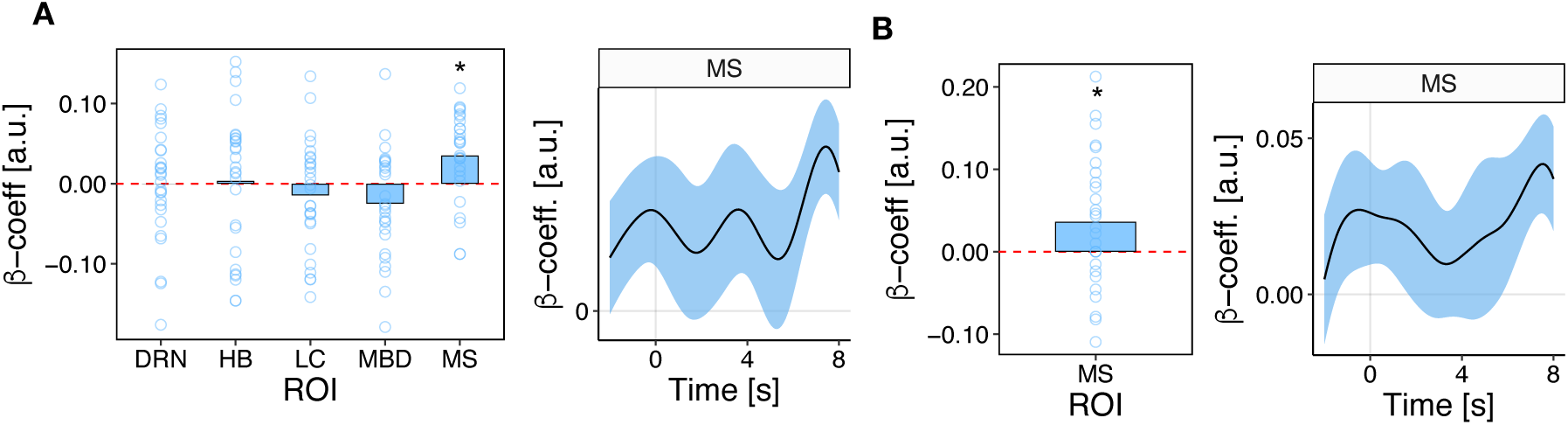
Activity in the medial septal nucleus represents changes in behavioural policy that are not driven by the environment. **(A)** BOLD signal in the MS ROI was modulated by incongruent behavioural policy switches – in other words, pursue-to-reject policy switches in poor environments and reject-to-pursue switches in rich environments *t_MS; incgongruent-switch_* (26)*=*3.35, *p*=.012. No other subcortical ROI signalled such events. **(B)** Follow-up analysis indicated that activity in MS signalled incongruent policy changes for the 10-point option specifically (*t_MS; incgongruent-switch 10-point_* (26)*=*2.32, *p*=.023). This is broadly consistent with the link between cholinergic pathways and explorative forms of behaviour, in the sense that incongruent policy switches involve deviation from an established reward-maximising strategy. In timecourse graphs, lines and shadings show mean ± SEM of β weights across participants. In effect-size graphs, bars show sample-mean magnitude of peak regression weights according to an unbiased leave-one-out procedure (see *Methods*). Dots indicate peak regression weights for individual participants.

